# Structure-Based TCR–pMHC Binding Prediction and Generalization to Unseen Peptides

**DOI:** 10.64898/2026.02.21.707231

**Authors:** A N M Nafiz Abeer, Raj S. Roy, Xiaoning Qian, Byung-Jun Yoon

**Author notes:** Contributing authors.

## Abstract

The interaction between T-cell receptors (TCRs) with the peptide-bound major histocompatibility complex (MHC) intricately impacts the functional specificity of T-cell-mediated adaptive immune response. Consequently, implication in immunotherapy has contributed to the ever-growing computational methods for TCR recognition, which have recently attracted structure-based approaches due to advancements in protein structure modeling. Despite access to structural information of the predicted binding interface, graph neural network (GNN)-based TCR-pMHC binding specificity classifiers tend to show poor accuracy for samples with unseen peptides. In this work, we comprehensively assess the potential factors that critically impact the generalization performance of classifiers trained with computationally predicted structures. Specifically, our experiments focus on analyzing the sensitivity of such predictors to the interaction features in the TCR-pMHC interface and the structural uncertainty. Building on the analysis, we demonstrate how the design of classifier architecture with auxiliary training objectives can improve the generalization performance to novel peptides not yet seen during model training. Overall, our work highlights the challenges of unseen peptide generalization from different perspectives of the GNN-based classifier paradigm, showcasing the strengths and weaknesses of the current state-of-the-art approaches in the generalization landscape.

## 1 Introduction

Insights from CD8^+^ T-cell mediated adaptive immune response, governed by the interaction of T-cell receptors (TCRs) with the peptide and the *α*1-*α*2 helices of major histocompatibility complex (MHC-I), are important for diverse immunotherapies against viral pathogens, cancer, and autoimmune diseases. Computational approaches have been proposed to learn the rules for TCR recognition of the peptide-bound MHC utilizing the data from different experimental assays. As the majority of the predictors have relied on the sequence information of peptides, MHCs, and TCRs [1–7], they cannot directly leverage the 3D local structural characteristics of TCRs interacting with the peptide bound in the central binding groove of MHC-I. Since the determination of the first TCR-pMHC complex structure [8], our knowledge base [9] regarding the structural aspects of TCR-pMHC interaction has been expanding with the growth in experimentally determined structural data [10]. This covers from the MHC restriction with peptide specificity for the TCRs [11] to the canonical docking, including structures with exceptions of these characteristics. Leveraging such information through deep learning techniques, such as graph neural networks (GNN), supported by development in computational techniques for protein structure modeling, can offer a promising pathway for predicting TCR-pMHC binding specificity, as observed in other structure-based protein property, protein-protein/ligand/DNA interaction prediction tasks [12–15].

In particular, structure-based efforts [16, 17] showed the potential of structural features for predicting the binding interaction between peptide and MHC. Given the cross-reactivity aspects of TCRs, and the limited number of TCR-pMHC samples across peptide-MHC pairs [10], the progress of such structure-based approaches for TCR-pMHC prediction is contingent upon the development of specialized computational tools for TCR-pMHC complex modeling. Consequently, the existing structure-based efforts for TCR-pMHC can be categorized into two branches based on the two intertwined tasks: the TCR-pMHC complex structure prediction and the binding activity prediction. For example, Bradley [18] developed AlphaFold-based TCR-pMHC complex structure prediction pipeline (TCRdock), providing improved structure prediction performance for TCR-pMHC than pretrained AlphaFold-Multimer model [19]. Furthermore, the confidence metric score, such as predicted aligned error (PAE) of the predicted TCR-pMHC complex from this specialized tool was shown to be effective in identifying the correct peptide within a pool of decoys. On the other hand, the authors in [20] moved the focus from predicted structure confidence to the docking quality (DockQ score [21]) of the predicted complex by AlphaFold-Multimer. Specifically, they trained a geometric vector perceptron (GVP)-based [22] graph neural network to estimate the DockQ score, which was applied in discriminating the binding and non-binding complex structures in a zero-shot manner. Complementary to both model confidence and docking quality, some research works [23–25] explored the application of interaction energy (estimated using tools like Rosetta [26], CHARMM [27] or contact-map derived pseudo-potential [25]) as a structure-based scoring function for TCR-pMHC binding. In this group of efforts, the binding activity prediction is considered as a zero-shot task where the performance relies on improvement in the structural modeling. The other branch focused on utilizing binding activity data in conjunction with the dataset of predicted complex structures. Slone et al. [28] proposed a GNN-based binary classifier for predicting TCR-pMHC binding activity from the residue level graph of TCR-pMHC complex structures predicted from the sequences. This is extended in [29] where the sequence encoding from ESM-2 [30, 31] was used to augment the node attribute of the TCR-pMHC residue graph, similar to LM-GVP framework [32]. In [33], specific data augmentation strategies, including re-docking and perturbing structures, were adopted to build the dataset of positive and negative samples from experimental TCR-pMHC structures. Such augmented dataset was used in training a hierarchical GNN classifier from residue and atomic level graphs, which showed reduced performance in TCR-pMHC binding classification when structures from computational tools, e.g., AlphaFold3 [34] and TCRmodel2 [35], were used instead of experimentally determined complexes.

With the advancement in computational protein complex structure prediction [19, 34, 36, 37], the increasing number of computationally predicted TCR-pMHC complex structures naturally drives the research effort for TCR-pMHC binding prediction toward a structure-based approach like GNN-based classifiers. While the existing works for GNN-based classifiers showed reasonable performance for test samples with unseen TCRs, these models performed very poorly for TCR-pMHC samples with novel peptides, i.e., unseen in the training dataset. One may speculate that such poor generalization performance is simply due to the quality of the training dataset, which can be improved by creating more (and better) samples. Additionally, it is also possible that the current computational dataset may have subtle features that are difficult to pick up for current models during training, and in such cases, increasing the dataset size may be marginally beneficial. As we move on to a larger training dataset scenario, it raises the questions like: “do such computationally predicted datasets have distinct characteristics required for generalization and accessible to our prediction models?” “Can we effectively leverage such *predicted* TCR-pMHC complex structures for improving the generalization performance for unseen peptides?” Exploring this line of questions is necessary for providing insights toward improved generalization performance for unseen peptides. Our work focused on this aspect of GNN-based classifiers for TCR-pMHC binding, underexplored in contrast to sequence-based approaches [38, 39].

Specifically, we assessed the unseen peptide generalization performance of GNN-based classifiers from three perspectives as illustrated in Figure 1: a) relative importance of interactions among peptides, MHC and TCRs in the computationally predicted structure in governing the generalized performance, b) influence of structural uncertainty in predicted TCR-pMHC complex structure on the binding prediction performance, c) sensitivity of generalization performance to the choice of classifier design. To enable balanced evaluation to explore these directions, we began with strict peptide splitting of the dataset from [28] without compromising the training dataset size, which reduced the bias in the results due to training sample size discrepancy. For assessing the impact of structural uncertainty, we generated two more datasets through different computational tools for the same TCR-pMHC samples, allowing inspection of the performance limitation imposed by the choices of such predicted datasets. Through highlighting the performance sensitivity of such GNN-based classifiers across different testsets, we demonstrated potential strategies, such as our proposed dual encoder approach [40] and training with preference data, for moving up the generalization performance further away from the random classifier-like performance. Altogether, our work shed light on the strengths and weaknesses of the structure-based TCR-pMHC binding prediction approach in tackling the challenges of unseen peptides.

**Fig. 1:**
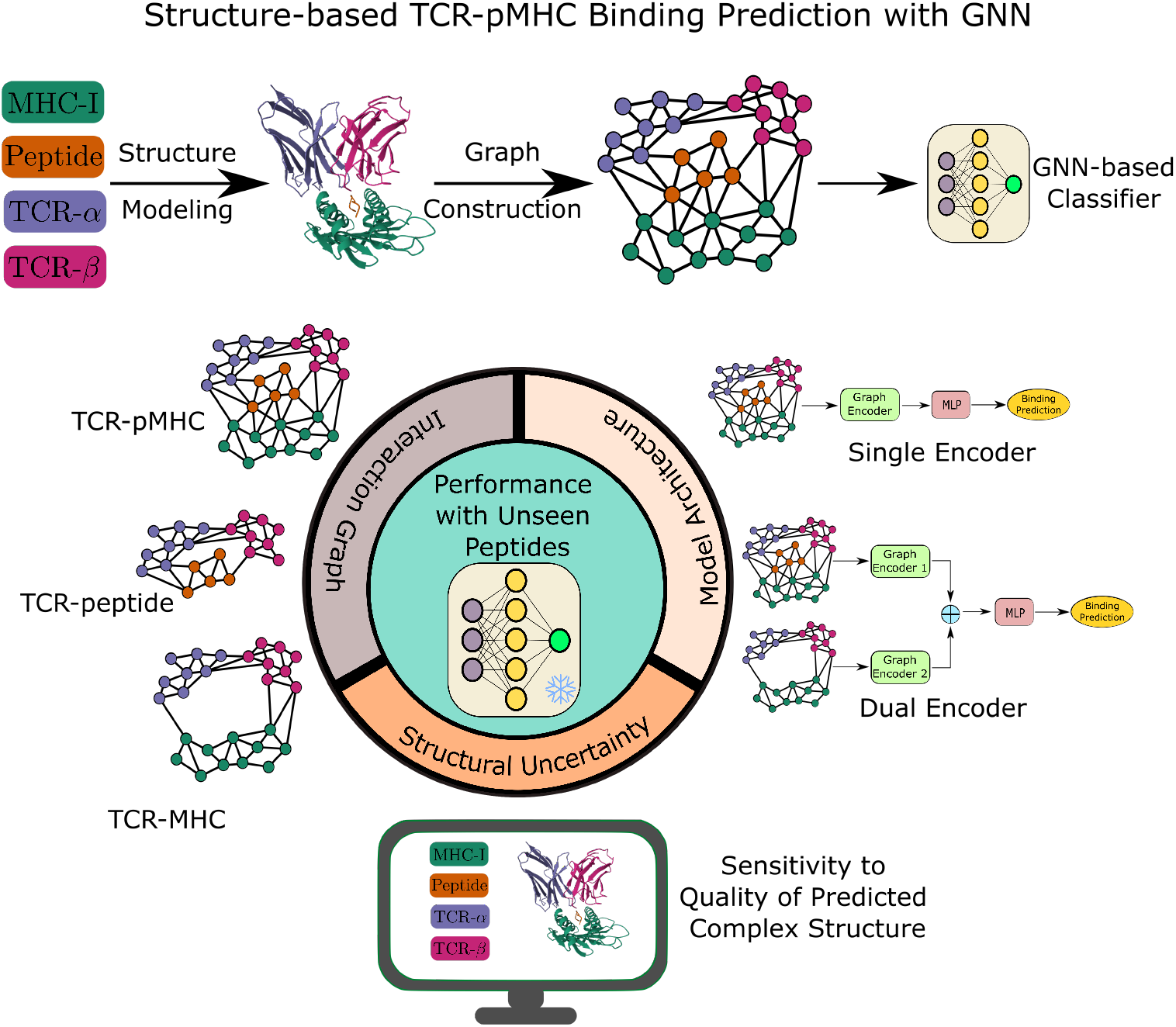
Overarching theme of our study: Structure-based TCR-pMHC binding prediction pipeline with graph neural network (GNN) and the potential factors (explored in this work) influencing the performance with unseen peptides. The GNN classifies a TCR-pMHC sample as binding or non-binding based on the residue-level graph constructed from the interface of predicted complex structure modeled by computational tools using MHC, peptide, and TCR-*α* and *β* sequences. Our study examines three aspects that can impact the performance of GNN-based models on novel peptides: the type of interaction edges, the quality of TCR-pMHC structure prediction, and the design of the GNN-based classifier.

## 2 Results

### 2.1 Evaluation with Peptides Unseen in Training

To evaluate the generalization performance of the structure-based approach, we considered the dataset of predicted TCR-pMHC structures from [28], comprising 345 unique peptides over 19282 TCR-pMHC samples. While the large pairwise normalized Levenshtein distance (in Figure 2**a**) among these peptides makes this dataset suitable for unseen peptide evaluation, the skewed TCR-pMHC sample distribution (Figure 2**b**) makes it difficult to make a balanced testset through random partitioning based on the peptide. Specifically, to fairly evaluate the performance of a prediction model across different testsets, one needs to keep the corresponding training datasets’ sizes similar. But due to the uneven number of samples per peptide, naive random splitting leads to testset of different sample sizes as well as peptide counts, and consequently different training dataset sizes. In our work, we constructed 5 different testset clusters from this dataset such that each contains equal number (7) of peptide samples, and approximately 1000 TCR-pMHC samples. We distributed 35 peptides (from the shaded region shown in Figure 2**b**) among 5 testset clusters in such a sequential manner (see Section 4) that each cluster has samples close to 1000. As illustrated in Figure 2**c** (in addition to contact map profiles in Figure A2), the interface geometries of binding TCR-pMHC samples in these clusters are distinct, offering a broad evaluation of the structure-based models.

**Fig. 2:**
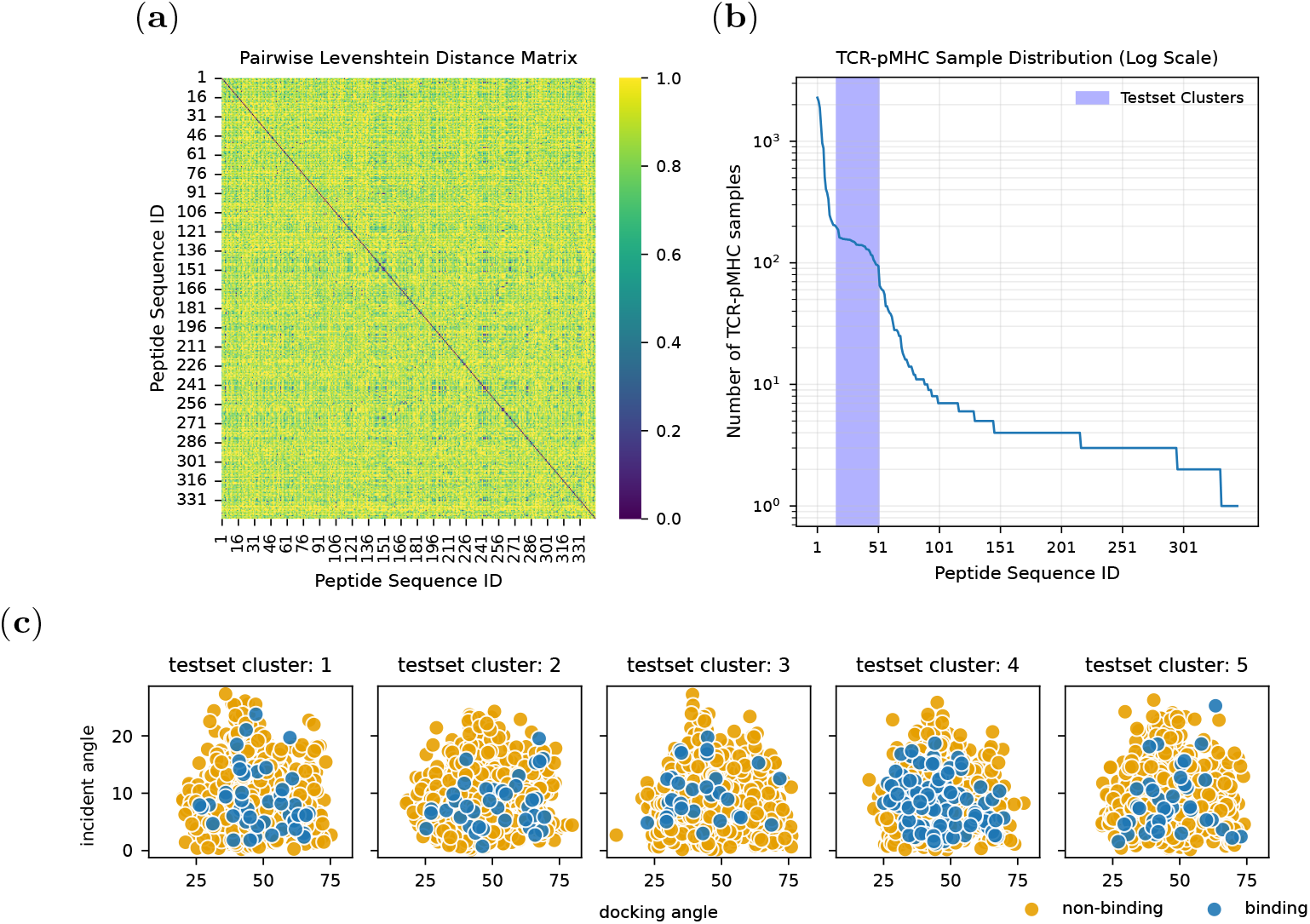
Peptide characteristics in dataset from [28] and binding geometry of the holdout testset clusters. (a) Pairwise Levenshtein edit-distance for the 345 unique peptides present in the dataset. (b) Distribution of TCR-pMHC samples where the shaded region indicates the testset peptides. (c) Docking and incident angles of predicted TCR-pMHC complex structures in testset clusters.

For evaluation on each of these 5 holdout testsets, we trained the GNN-based classifiers with the remaining data (removing samples of the holdout testset from the whole dataset of 19282 samples) in 10-fold cross-validation manner, and performed prediction for the holdout test samples with 10 trained models (corresponding to the 10 folds). We considered two types of graph encoder: Equivariant Graph Neural Network (EGNN) [41] and Edge-Variable-Transformer convolution (EVTConv) [28] in single and dual encoder-based classifiers [40] (see Section 4 for details). In our proposed dual encoder approach, both the TCR-pMHC interface residue-level graph and its TCR-MHC subgraph are encoded in such a manner that encourages these two embeddings to be aligned or different depending on the training label (binding and non-binding) through a semantic loss term combined with binary cross-entropy loss. Note single encoder approach with EVTConv as graph encoder is identical to the classifier proposed in Slone et al. [28].

As shown in Table 1 and Figure 3 (boxplots without the hatches), there is no clear winner among these 4 classifiers that generalizes to all 5 testset clusters. Among the graph encoders, the EGNN showed more robustness (higher median) compared to EVTConv. In terms of classifier architecture, the dual approach sometimes out-performed the single encoder counterpart (cluster 1, 2 and 3), and such trend showed sensitivity to the choice of graph encoder. Note this trend of the dual encoder approach is influenced by its semantic loss component (see Figure B3, Section B). That being said, this improvement over the single encoder approach illustrates that the dataset of predicted structures has a certain amount of relevant signal useful for improving the generalization performance, but requires efforts in designing a robust classifier architecture. Interestingly, none of the four classifiers was able to significantly move up from the random classifier baseline (i.e., AUC-ROC of 0.5) for testset cluster 5, illus-trating the need for carefully designed unseen peptide splitting-based evaluation for benchmarking GNN-based approaches in TCR-pMHC binding prediction.

**Table 1:** ROC-AUC performance for 5 holdout test clusters. For Boltz-2 [36] and FoldX-5.1/ Rosetta [26, 42, 43], the confidence measures and (negative of) interaction energies (between TCR and pMHC), respectively, are used for classification of binding and non-binding samples. For GNN-based approaches, a 10-fold cross-validation training was first performed on the training dataset, and the resulting 10 models were then applied to the holdout testsets. We reported the median and interquartile range (IQR) over the 10 ROC-AUC values.

| Method | Cluster 1 | Cluster 2 | Cluster 3 | Cluster 4 | Cluster 5 |
| --- | --- | --- | --- | --- | --- |
| Boltz-2 iptm | 0.5707 | 0.5574 | 0.6022 | 0.6263 | 0.5636 |
| Boltz-2 confidence score<br>($0.8 \times \text{pLDDT} + 0.2 \times \text{iptm}$) | 0.5742 | 0.6480 | 0.5547 | 0.5997 | 0.5757 |
| FoldX-5.1 interaction energy | 0.4367 | 0.5255 | 0.3963 | 0.5355 | 0.4858 |
| FoldX-5.1 interaction energy<br>(backbone only) | 0.5069 | 0.4941 | 0.4790 | 0.5902 | 0.5705 |
| Rosetta interaction energy | 0.5368 | 0.5545 | 0.5707 | 0.5495 | 0.5276 |
| EGNN (single) | 0.5548 (0.0372) | 0.6108 (0.0750) | 0.5506 (0.0658) | 0.6718 (0.0454) | 0.5239 (0.0613) |
| EVTConv (single) | 0.5569 (0.0639) | 0.5346 (0.0524) | 0.5403 (0.0686) | 0.6195 (0.1228) | 0.5072 (0.0678) |
| EGNN (dual) | 0.6012 (0.0607) | 0.5899 (0.0448) | 0.5760 (0.0491) | 0.6045 (0.0793) | 0.5002 (0.0811) |
| EVTConv (dual) | 0.5453 (0.0688) | 0.5899 (0.0802) | 0.5532 (0.0902) | 0.5976 (0.1007) | 0.5322 (0.0751) |

**Fig. 3:**
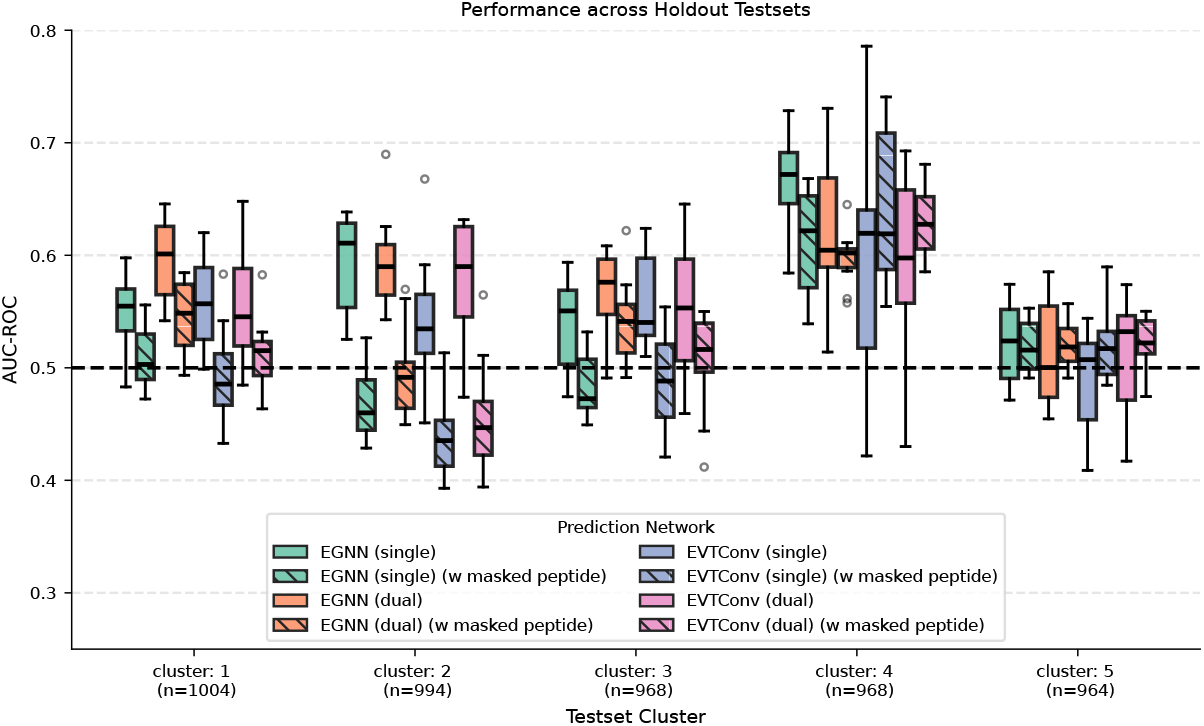
Performance of 4 different GNN-based classifiers with inference on complete TCR-pMHC interaction graphs and the subgraph with peptide nodes masked. These four models (two types of graph encoder: EGNN and EVT-Conv, and two types of classifier architecture: single and dual encoder approach) are trained with complete TCR-pMHC graphs. The horizontal dashed line at 0.5 refers to the performance of a random classifier.

The performance of “zero-shot” baselines (Table 1) further shows the utility of the GNN-based approach with predicted structures. All GNN-based classifiers exceeded the interaction-energy-based screening in terms of median performance, except for cluster 5, where the FoldX-estimated energy was better. However, compared to Rosetta-derived energy, the performance of FoldX was highly inconsistent across different testset clusters. On the contrary, the confidence metric-based approach with Boltz-2 showed more balanced performance, and the training of GNN-based classifiers with TCR-pMHC structures led to better (median) classification performance over Boltz-2 only in clusters 1 and 4.

### 2.2 Influence of peptide in structure-based prediction

As discussed in Section 2.1, the performance of GNN-based classifiers was improved for some clusters with the dual encoder approach, which utilizes TCR-MHC and TCR-pMHC interface graphs. Since the peptide nodes are in a small fraction of the total nodes (with a lower fraction of edges involving peptide, as shown in Section A) in the TCR-pMHC interface graph, we investigated whether they are crucial for the prediction performance or the dual encoder approach was simply bypassing them with the TCR-MHC subgraph. We repeated inference of 4 classifiers considered in Section 2.1 with the same testset clusters, but this time the TCR-MHC graph (instead of TCR-pMHC) is passed to the GNN. Figure 3 revealed three distinct characteristics of the GNN-based prediction. Firstly, the masking of peptides led to a significant drop in performance in clusters 1, 2, and 3, indicating that the peptides played a role in performance improvement in these clusters by the dual encoder approach. In cluster 4, we observed a sensitivity to the choice of graph encoder. Interestingly, the performance with EVTConv improved slightly when we performed inference by removing the peptide nodes. This behavior is quite the opposite of EGNN, which showed a diminishing performance. Finally, there is no significant change in performance for cluster 5, implying the prediction of GNNs on this cluster is not heavily impacted by the peptides, which potentially explains why the generalization performance for this cluster significantly differs from that of other testset clusters.

#### Utility in interpretability

For a model trained with TCR-pMHC structures, the learned classification rules ideally should involve the edges involving the peptide nodes in the graph. Intuitively, such model’s confidence in its prediction for a test sample is expected to decrease when the peptide nodes are removed from the graph. For instance, if a model predicts a test sample’s TCR-pMHC residue graph to have a chance of 80% to be binding, and increases the chance to 90% for the same sample with peptides removed, the earlier prediction of 80% seems less reasonable than it would have been if the prediction decreased toward 50%. This change in predictive entropy due to peptide masking can be used to pick the “best” prediction from 10 models corresponding to 10-fold cross-validation. We discussed such approaches in Section C.

### 2.3 TCR-pMHC vs TCR-MHC vs TCR-peptide: What matters more?

The variation in performance due to the masking of peptides in test time (Section 2.2) led us to the next question: what interactions among peptide, MHC, and TCRs are crucial for improved generalization performance? We trained classifiers with the TCR-MHC and TCR-peptide interface residue level graphs by masking peptide and MHC nodes, respectively. With two separate GNN encoders, we have four new classifiers to compare (Figure 4) against the single and dual encoder approaches, which start from the complete interface graph.

**Fig. 4:**
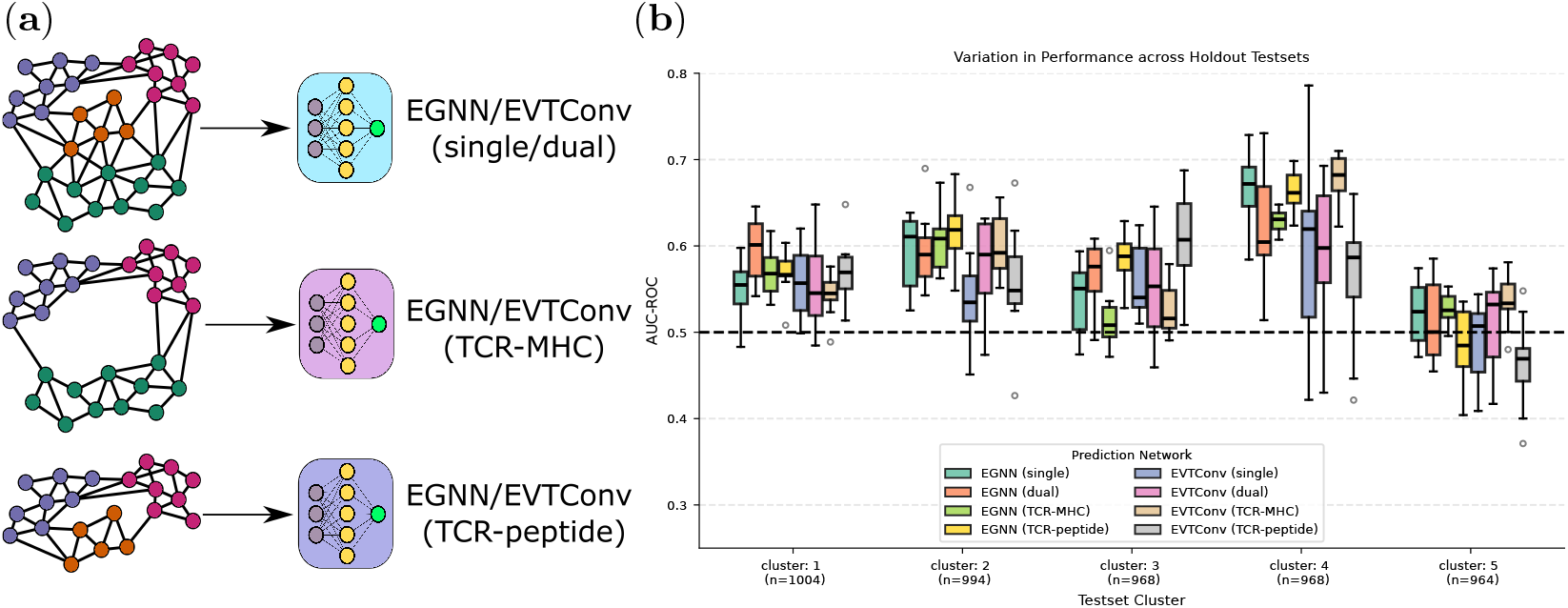
Performance of models trained (and tested) with TCR-pMHC, TCR-MHC, and TCR-peptide (interface) residue level graphs.

#### TCR-MHC

Let’s first focus on clusters 1, 2 and 3, where the dual encoder showed performance improvement over the single encoder approach. For cluster 1 and 2, TCR-MHC seemed to be enough as the models trained with TCR-MHC showed similar performance with the dual encoder counterpart. However, as seen in Section 2.2, the performance of dual encoder approaches is sensitive to the peptide nodes, which demonstrates that the dual encoder approach is not overlooking the peptides in preference of the TCR-MHC interface graph, even though we have a second encoder that directly encodes the TCR-MHC graph in this approach. This point is further strengthened by the decline in performance in cluster 3 with the models trained with TCR-MHC. Specifically, the generalization performance drops below that of the single and dual encoder approaches, indicating that the TCR-MHC is not as important for this cluster as it is for the first two clusters. This illustrates how the dual encoder approach balances the influence of different interactions during training of the classifier. Now looking back at the result of cluster 4 with peptide masking during inference time (Section 2.2), we saw improvement in generalization performance with EVTConv. This observation of peptides negatively influencing the performance of EVTConv is further supported by the improvement in AUC-ROC of cluster 4 when we trained the models (with EVTConv as graph encoder) with the TCR-MHC only graphs. Now this also potentially explains why the dual encoder approach performs poorly compared to the single encoder approach, since the former one tries to make the interaction related to peptides more influential in its decision-making.

#### TCR-peptide

Performance drop in cluster 5 with models trained with the TCR-peptide graph through masking of the MHC nodes demonstrates the MHC being the “useful” (albeit for the performance similar to the random classifier) source of information learned by the GNN-classifiers for this cluster. Combining with the observation that generalization slightly improved (EVTConv) or stayed similar (EGNN) for models trained with TCR-MHC, it explains why this cluster showed no significant change in the performance of different GNN-classifiers in the peptides masking experiment of Section 2.2. For cluster 4, the performance of TCR-peptide-only models is significantly worse than the TCR-MHC-only models for EVTConv as the graph encoder, while for the case of EGNN, the scenario is opposite. Furthermore, ignoring MHC during training led to improved generalization performance (compared to the single encoder approach that uses all node types) in clusters 1, 2, and 3, irrespective of the graph encoder type. For example, in the case of cluster 3, the median AUC-ROC of EVTConv classifiers trained with peptide-TCR exceeded 0.6, whereas the corresponding classifiers (single encoder) trained with TCR-pMHC have a median performance of around 0.54. Since the dataset has the least diversity in terms of MHC, it is intuitive to expect that ignoring the information from MHC during training classifiers may not hurt the performance of classifiers. While the results from the first three testset clusters are aligned with this idea, the case of clusters 4 and 5 showcased the situation where the signals from MHC could be significant for better generalization performance depending on the types of graph encoders employed in the classifiers.

Overall, the discrepancy in performance across testset clusters associated with the choice of both graph data and graph encoders highlights the intricate learning rules extracted in the training process. Not all interactions among TCRs, peptides and MHC may be uniformly present (Figure A1 and Section A) in the testset, so betting on a particular interaction during training can hurt the performance by losing other information. We speculate that classifiers that allow the learning of classification rules by balancing the importance of these interactions are better suited for generalizable performance. Dual encoder (with EGNN in particular) is along this direction as demonstrated in Section 2.4, but is still biased (by design) to the interactions associated with the peptide.

### 2.4 Impact of varying degree of utilization of TCR-MHC context

During inference for TCR-pMHC samples with an unseen peptide, the TCRs (and MHC) of the test sample can already be present in the training dataset. Having a separate encoder for TCR-MHC can be thought of as a way for encoding this “context” representation for the peptide in the TCR-pMHC sample. With the dual encoder approach, the encoding of the TCR-pMHC is only implicitly influenced by the encoding of TCR-MHC through the combined loss of binary cross-entropy and the semantic loss. In this section, we analyzed what if we allowed the TCR-MHC encoding to have a more dynamic impact on the encoding of the TCR-pMHC graph?

Accordingly, we considered two approaches (see Section 4) to perform conditional encoding of the TCR-pMHC graph with the subgraph TCR-MHC as a condition. In the first alternative, we fed the graph-level representation of the context (TCR-MHC) as a global node feature to the encoding stage of the TCR-pMHC graph. The second alternative involves injecting the context into the encoding of TCR-pMHC graph through the parameters of its node embedding MLP layers. Specifically, the parameters of the node embedding MLP layer are predicted by the graph-level representation of the context of each sample. From Figure 5, we see that the dual encoder approach has a balanced performance between these two levels of utilization for clusters 1-3. On the other hand, the performance deteriorates for clusters 4-5 as we make the dynamic encoding stronger. Interestingly, the dual encoder approach without the semantic loss component (see Section B for details) also showed a decline in performance similar to that of the first alternative (i.e., context as global node feature)–illustrating the importance of semantic loss formulation. These results re-emphasize the intuition that over-reliance on a particular interaction (TCR-MHC in this case) can weaken the robustness of the classifiers.

**Fig. 5:**
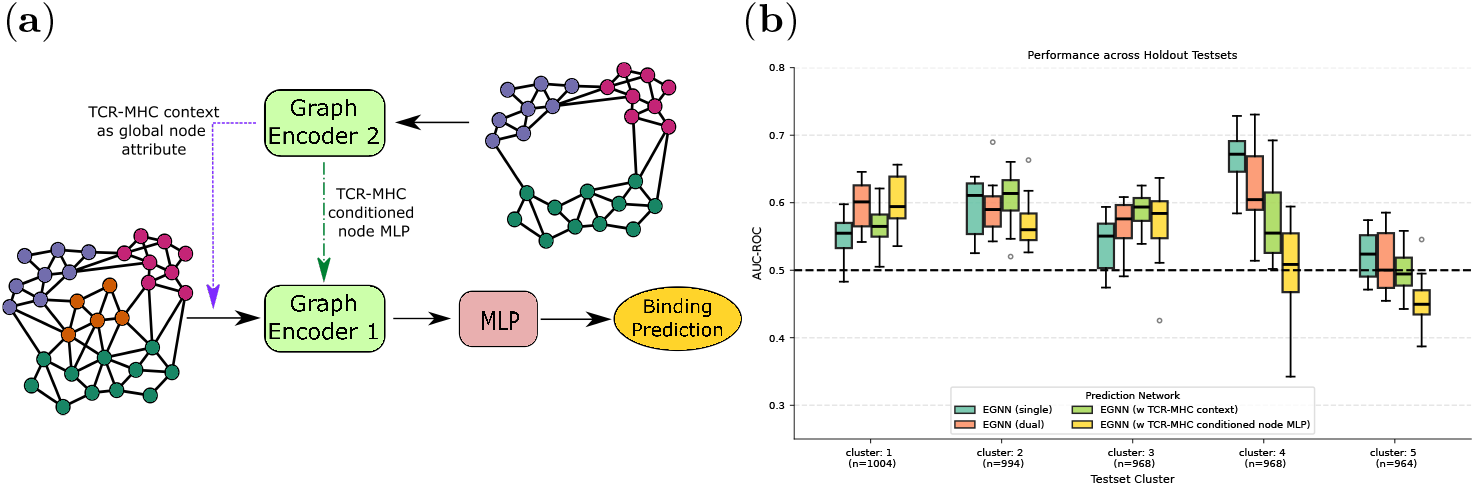
Impact of varying levels of utilization of TCR-MHC context. (a) The encoded representation of TCR-MHC residue-level graphs is either used as a global node attribute or as a condition to generate the parameters of node embedding MLP in the encoding of the TCR-pMHC interaction graph. (b) Unseen peptide performance comparison with single and dual encoder approaches with EGNN as graph encoder.

### 2.5 Influence of structural quality for generalization

Since we are using predicted TCR-pMHC complex structures in the GNN-based classifiers, the generalization performance is inherently impacted by the structural quality of the predicted complex. To analyze the sensitivity to the predicted structures, we constructed two new datasets based on the dataset from [28]. The first one (denoted as *relaxed struct*) consists of Rosetta-relaxed structures where the samples from [28] are used as initial structures for relaxation. For the other dataset (denoted as *tcr-model2 struct*), we predicted the TCR-pMHC structures from the same sequences using TCRmodel2 [35]. We then repeated the evaluation of Section 2.1 with these two new datasets where we kept the same hyperparameters for the classifiers.

Figure 6 depicts the holdout test results with *relaxed struct* dataset. In terms of root mean square deviation (RMSD) between initial and relaxed structures, there are not many noticeable patterns across the testset clusters, except that a few binding samples of cluster 5 deviated more due to relaxation (Figure 6**a**). In terms of generalization performance, we observed a downward shift (either median or 3^rd^ quartile decreased) in AUC-ROC for all classifiers in the case of cluster 5. For other testset clusters, most of the models showed similar performance (similar to models trained with initial structures) with a couple of exceptions. For instance, both single encoder-based classifiers observed a boost in performance for cluster 3.

**Fig. 6:**
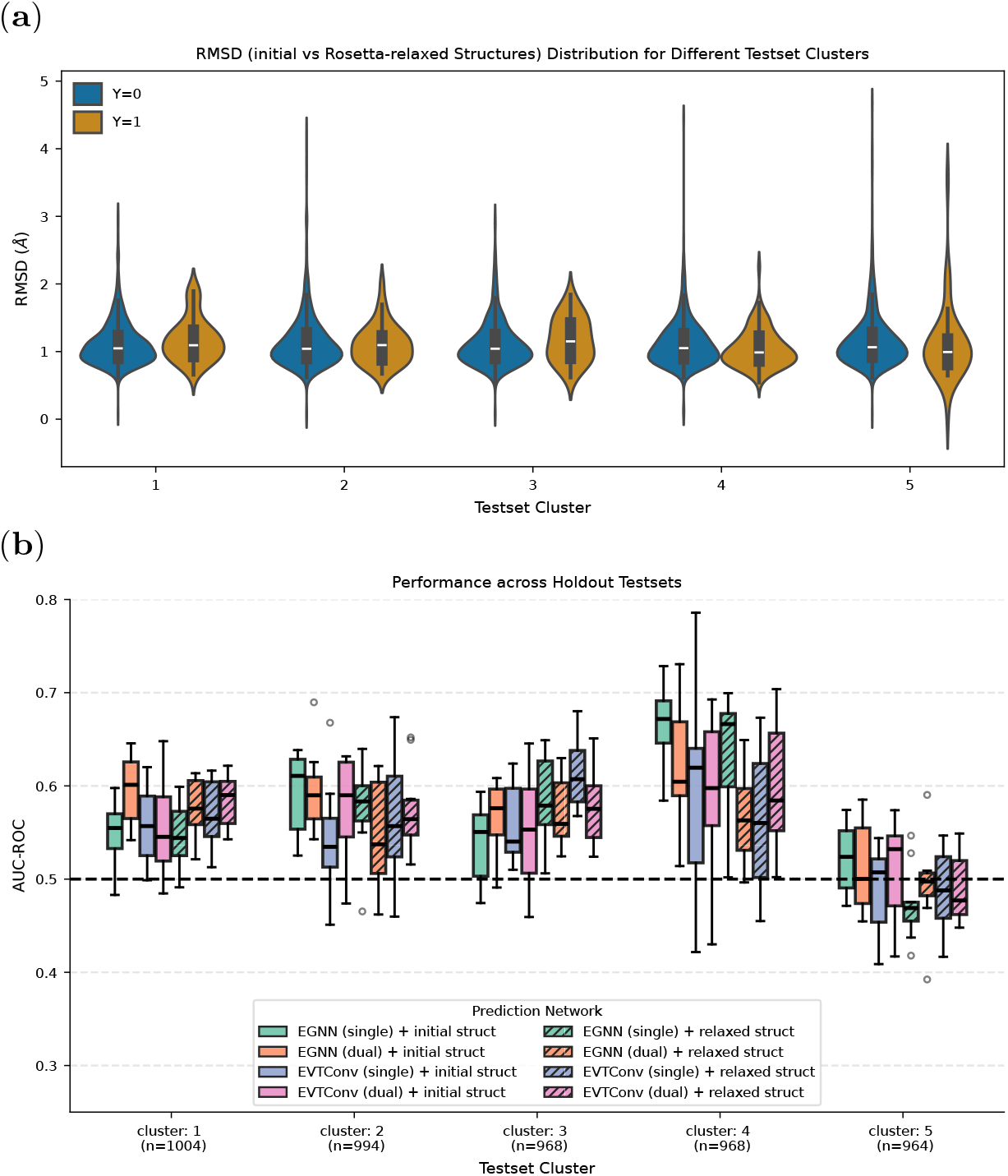
Impact of Rosetta-relaxation of TCR-pMHC structures. (a) Class-specific violin plots of RMSD (Å) between relaxed and initial structures in each holdout testset. (b) 4 different classifiers trained with relaxed structures are compared against the corresponding classifiers trained with initial structures.

Considering the fact that relaxation might have distorted the structural artifact (which the GNN can exploit) in the predicted structures, such variation in testset performance due to datasets of different structural quality raises the question of which predicted structures to use for training the GNN-based classifiers. Experiments with *struct tcrmodel2* showed similar variation in performance (see Section D), including the challenging scenario of testset cluster 5.

### 2.6 Avenues for improving the generalization performance

The focus of this section is to briefly discuss empirical efforts to improve the generalization performance on the testset cluster 5 (see Section E for efforts based on training-data dependency). Previously, we observed that the MHC nodes are important for the different classifiers’ performance in cluster 5. So, we investigated whether explicit incorporation of human leukocyte antigen (HLA) type information of MHC molecules can help in improving the performance. Accordingly, we trained a variant of the dual encoder approach where the one-hot encoding vector of HLA type is concatenated with the encoded graph representations of TCR-pMHC and TCR-MHC before passing to the MLP layers for final binding probability prediction.

Additionally, we also observed (Table 1) that the Rosetta-energy-based scoring mechanism [23] showed somewhat stable performance (across testset clusters) slightly above the random classifier. We wondered whether the biophysical energy data of the predicted TCR-pMHC complex could be leveraged in a way that complements the typical binary label-based training paradigm. Along this direction, from each fold’s training datasplit 𝒟_train_, we created a paired preference data 𝒟^pair^ = {(𝒢 _*i*_, 𝒢 _*j*_)|*y*_*i*_ = 1, *y*_*j*_ = 0, Δ*E*_*ij*_ ≤ δ_th_}, where each paired sample consists of positve and negative samples with same peptide, MHC sequence, but different TCRs. Δ*E*_*ij*_ is the difference between positive and negative samples in terms of TCR-pMHC interaction energy estimated by Rosetta, and we chose to include those samples with a negative energy difference, i.e., lower energy for binding samples. See Section 4 for the details of this construction procedure. In addition to training with binary class labels from 𝒟_train_, we periodically trained the classifiers with 𝒟^pair^ to penalize predicting a lower probability for the positive samples than the negative ones with shared pMHC.

Figure 7 demonstrates the impact of these two approaches for cluster 5. Use of only HLA, makes the performance of the dual approach more consistent (lower IQR) with slightly better median AUC-ROC. On the other hand, the periodic training with preference data boosted the performance of both single and dual encoder approaches, i.e., an increase in median. While such improvement is still close to the performance of a random classifier, this upward trend indicates the utility of biophysical energy estimates in enabling the GNN-based classifiers to extract the features that are rarely reached through the binary label-based training. However, this preference loss-based training comes with the disadvantage of introducing sample-specific bias. Specifically, 𝒟^pair^ only includes pairs of positive and negative samples which follow Rosetta-derived energy relation (i.e., positive samples to have lower energy than the negative one in the pair). To reduce the deviation from the classifier trained with binary cross-entropy loss, the periodic training (with a lower learning rate) strategy is employed in this work. As we showed in Section F, this strategy can, in some cases, fail to find better models than trivial binary classification-based training. Nonetheless, such approach with interaction energy estimates seems to be one of the potential directions to crack the challenging cases like cluster 5.

**Fig. 7:**
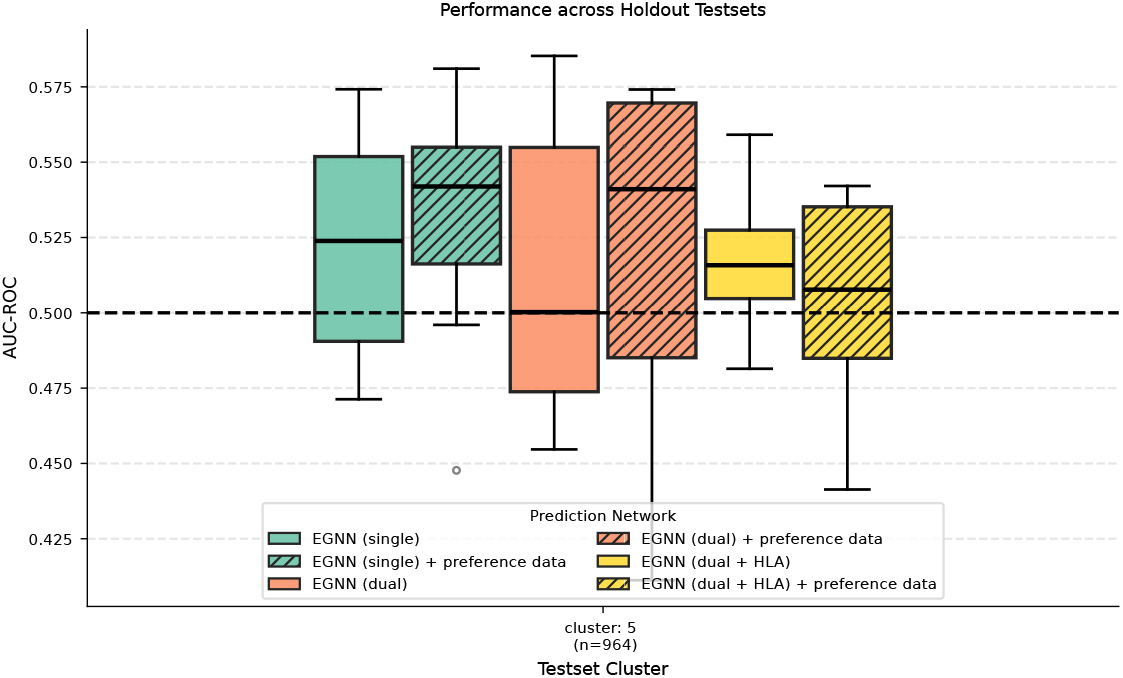
Impact of training with paired preference data. The single and dual encoder-based classifier is same as in Section 2.1, and the third model denoted as “dual + HLA” has one-hot encoding vector of HLA type concatenated with encoded graph representations of TCR-pMHC and TCR-MHC. In case of preference data training, the models are trained periodically with a dataset of 3000 preference pairs (from training samples in the corresponding fold) in addition to the usual per-epoch training process.

## 3 Discussion

Learning from the structural characteristics of TCR-pMHC interaction is an intuitively promising direction to address the concerning issue of generalization performance of current deep learning predictors in cases of unseen peptides. In this work, we explored the potential of the graph neural network-based classifiers in improving the scenario by leveraging the dataset of computationally predicted structures of TCR-pMHC complexes. Our experiments with different holdout testsets revealed the sensitivity of the generalization performance to the choice of graph encoder as well as the relative importance of specific interactions across different testsets. For example, the same neighborhood context of the peptide, i.e., TCR-MHC interface graph is as informative as the TCR-pMHC interface in some test clusters, but faces difficulties in identifying binding and non-binding samples in other test cluster. Moreover, in contrast to the sequence-based approaches that often consider only the pairs of peptide and TCRs, our results indicated mixed utility of MHC in structure-based TCR-pMHC binding prediction. Specifically, the large chain of MHC can be beneficial for test samples where key interactions in the TCR-pMHC graph learned by the classifiers are subtle, while in other clusters it can obscure the peptide-relevant information. Through different level of TCR-MHC utilization in GNN-based classifiers, we showed the robustness of our proposed dual encoder approach to balance the importance of all interactions in the TCR-pMHC interface, resulting in improvement with EGNN over single encoder approach which classifies the sample from encoded representation of only the TCR-pMHC interface graph. However, neither these efforts from the classifier architecture perspective were enough for improved generalization in a particular testset (cluster 5), nor did the dataset of computationally determined structures of different quality change the scenario. For this case, our periodic finetuning strategy with pairwise preference dataset based on Rosetta-estimated TCR-pMHC interface energy resulted in slight improvement, illustrating the need for auxiliary training objectives to enable the classifiers to focus on subtle characteristics of the binding interface that can be overlooked in binding specificity-based training.

In our work, the computational TCR-pMHC complex structure prediction pipelines are decoupled from the binding activity prediction network. While finetuning the structure predictor can be explored [16], a more relevant direction is quantifying the quality of a predicted structure based on not only the structure itself, but how it influences the classifier’s performance. Specifically, the latter approach can be used to prioritize the structural data to reduce the classifiers’ sensitivity to the choice of computational modeling tools. The original structural dataset from [28], and the two versions: *relaxed struct* and *tcrmodel2 struct* that we generated in our work, are relevant resources for pursuing this direction.

The list of potential factors impacting the generalization performance considered in our work is not exhaustive. In fact, the interaction between TCRs and the pMHC is dynamic in nature, that is difficult for GNN to learn from the static structures of the complexes. For example, the binding does not necessarily lead to functional activation of T-cell response [44] without the potential roles of other factors, such as the formation of a flexible catch-bond [45] at the TCR-pMHC interface. While insights from molecular dynamics (MD) simulation [46] can uncover unique characteristics of such complex interactions, integration of such knowledge into the graph encoding network is not straightforward. The development of the large structural datasets of TCR structures [47, 48] potentially takes the current deep learning approaches one step closer toward considering the flexibility of complementarity determining regions (CDR) loops of both *α* and *β* chains of T-cell receptors into the binding specificity prediction. Despite advancement in accelerated techniques for protein conformation generation [49], their incorporation into TCR-pMHC structure modeling warrants detailed investigation, as findings from recent studies [50] are not reassuring for protein-protein docking from the ensemble of conformations.

Our experiments demonstrated that the set of computationally predicted TCR-pMHC complex structures has different interaction characteristics that the graph neural network can lock onto as a shortcut during training, which do not necessarily lead to more generalized rules for discriminating binding samples with unseen peptides from non-binding ones. Through enforcing the classifier on particular interactions, models trained with the same computational dataset can show improved generalization performance, as illustrated with our proposed dual encoder approach. However, even for a cross-reactive TCR, the binding interfaces do not necessarily conform to the same 3D structural shape for different peptides [51]. Similar to the protein language model for sequence representation, the learned representation of the molecular complex with large-scale pre-training [13] can potentially help with such discrepancy between training and test samples in terms of structural aspects of TCR-pMHC. For instance, a pre-trained graph encoder in [52] was shown to be a reliable zero-shot predictor of binding affinity between TCR and a small group of antigens. Moreover, as the computational approaches [53, 54] for TCR-mimicking binder design targeting peptide-MHC complex continue to develop, the interaction profiles between the binder and the pMHC are likely to add new structural insights. While these designed binders may not have the same flexibility as TCRs, the insights can elucidate the local neighborhood preference near the pMHC interface for the binding activity.

Graph neural networks have been widely adopted for capturing the structural characteristics of intermolecular interactions, e.g., protein-protein complexes. Our work assessed the reality of its potential in the challenging task of predicting TCR-pMHC binding specificity for unseen peptides, where the generalization issue is exacerbated by the diverse binding mechanisms with a limited number of experimentally determined structures. We hope that our computational experimental findings with different GNN-based classifiers as well as the generated datasets, may facilitate further efforts to improve the prediction accuracy of the structure-based approaches with predicted TCR-pMHC structures.

## 4 Methods

### 4.1 Dataset and holdout testset selection

We collected the dataset of [28] (denoted as *init struct* in our work) from https://github.com/KavrakiLab/STAG_public, where 19282 TCR-pMHC structures are reported with the binary class labels of binding and non-binding samples. The authors of [28] curated the positive samples from multiple databases: McPAS-TCR [55], VDJdb [56], IEDB [57] and 10x genomics [58], while the TCR-swapping strategy was employed to augment the pool of negative samples in addition to samples from 10x genomics. The TCR-pMHC structures were modeled with TCRpMHCmodels [59] and ImmuneScape [60] based on the sequences of peptide, MHC and TCRs (*α* and *β* chains obtained via Stitchr [61]). The details of this data curation process can be found in [28].

Building on top of this dataset, we further created two additional datasets: *relaxed struct* and *tcrmodel2 struct*. Specifically, *relaxed struct* comprises Rosetta relaxed structures of all samples from *init struct*. For the latter case, we predicted the structures with TCRmodel2, which can sometimes end up predicting structures with unrealistic binding or have low prediction confidence. Since TCRmodel2 generates 5 predicted structures for each TCR-pMHC sample, we retained the prediction with the highest model confidence (exceeding 0.49, based on the recommendation in https://tcrmodel.ibbr.umd.edu), which includes inter-chain edges between the peptide and TCRs. Consequently, we had to remove 987 samples (out of 19282), which includes 117 binding samples. Note that this set of removed samples includes one case where the TCRmodel2 program (both local and online server) failed.

#### 4.1.1 Holdout testset construction

We first rearranged the 345 unique peptides from *init struct* according to the counts of their corresponding TCR-pMHC samples. Then we picked the shaded region (Figure 2**b**) of 35 peptides since they have a more uniform sample distribution while allowing the testset size to be around 1000. To minimize the sample discrepancy among testset clusters, we first assigned the top 5 peptides (within these 35 selected peptides) to clusters 1, 2, 3, 4, and 5, respectively. For the assignment of the next top 5 peptides, we reversed the order, i.e., cluster 5 to cluster 1. This assignment process was repeated until all 35 peptides were distributed to these 5 holdout testset clusters. Note, this sequential distribution strategy with periodic reversal of assignment order was applied to make sure that GNN-classifiers utilized a similar number of training samples for each holdout scenario, which allowed us to analyze their performance discrepancy across different testsets.

### 4.2 Structure-based binding prediction

#### 4.2.1 Zero-shot baselines

##### Boltz-2

For each holdout testset, we predicted the TCR-pMHC complex structures with Boltz-2 [36] from the sequences, and used the interface predicted template modeling (ipTM) score and the Boltz-2’s overall confidence score (0.8 *×* pLDDT + 0.2 *×* ipTM) to discriminate the binding vs non-binding samples. Here, pLDDT is the predicted local distance difference test for the predicted structure.

##### Interaction energy-based

We computed the interaction energy between pMHC and the *α* and *β* chains of TCRs in the predicted TCR-pMHC complex structures of *init struct* dataset. FoldX-5.1 [42] and Rosetta [26, 43] are used to estimate the biophysical energy between these two groups (pMHC and TCRs), preceded by each tool’s recommended pre-processing stages, i.e., repair and relax, respectively. The negative of such estimated energy is considered to rank the test samples, assuming that the binding samples have lower interaction energy than the non-binding cases.

#### 4.2.2 Graph neural network (GNN)-based classifiers

##### TCR-pMHC interface residue graph

Following the residue graph construction process of [28], we first identified the 3D coordinates of TCR-pMHC interface residues (*C*-*α* atoms) where the residue group of TCRs and the residue group of pMHC are within 14Å of each other. From the interface residues, the TCR-pMHC interface residue level graph (𝒢_TCR-pMHC_) is constructed, where edges exist up to an inter-residue distance of 8Å. For a fair comparison with [28], which is the single encoder with EVTConv in our framework, we used the same node and edge features for the residue level graph from that work. Specifically, the biophysical attributes of amino acids like Atchley [62] and Kidera [63], along with one-hot encoding of chain types, were used as the node features (***x***_feat_) of the TCR-pMHC interface graph.

For edge features, the radial basis function of distance (Equation (1)) with 15 different scales is concatenated with the edge type representing vector (Equation (2)). Here, ***τ***_*i*_ is a one-hot encoding vector to indicate the chain of residue *i*.

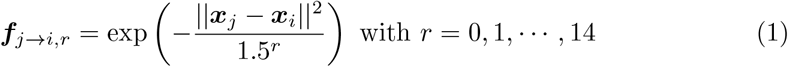

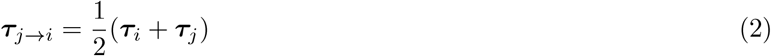

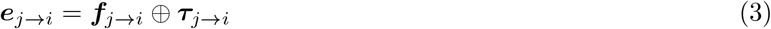

##### Graph encoder

To extract graph level representation, i.e., ***g*** = *ϕ*(𝒢), we considered two different graph neural networks: EGNN [41] and EVTConv [28]. The message passing steps for EGNN depicted in Equations (4) and (5) are different compared to [41] since we kept the ***x***_*i*,coords_ fixed during message propagation across *L* layers. The node feature ***x***_*i*,feat_ is projected to 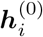 through a MLP block, *ϕ* _node-MLP._ *ϕ*_*e*_ and *ϕ*_*h*_ are MLP blocks for aggregating the messages from neighbors and combining the aggregated message with the current node embedding, respectively.

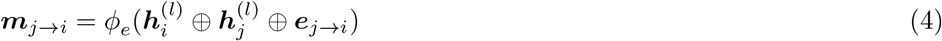

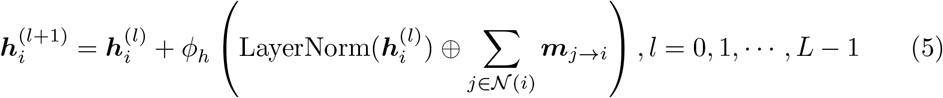

On the other hand, EVTConv has edge-specific attention score (*α*_*ij*_) to perform weighted message aggregation with edge-conditional encoding of the upcoming message via ***W***_*j*→*i*_. In Equations (6) to (8), ***W***_0_, ***W***_*q*_, ***W***_*k*_ are learnable weight matrices, and ***W***_*j*→*i*_ is predicted from edge embedding by a learnable MLP block *ϕ*_***W***_. Note the edge embedding 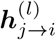 is encoded from 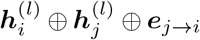 through another MLP block (different than the *ϕ*_*e*_ of EGNN).

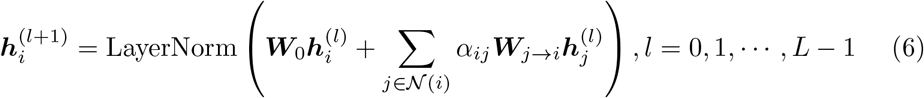

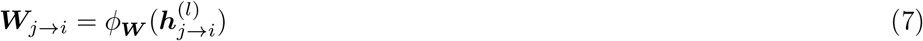

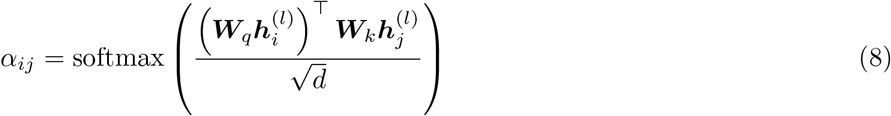

Finally, the graph level representation is obtained via mean-pooling (Equation (9)) of the updated node embeddings after *L* layers of message passing. We used *L* = 3 layers of GNN encoder with a message dimension of 32.

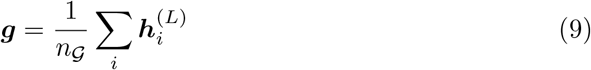

##### GNN-based classifiers

Typically, a residue-level graph is encoded by a graph neural network to produce a graph-level representation, which is further processed via a multilayer perceptron (MLP) block to predict the binding probability of the TCR-pMHC sample. We denote this approach as the *single encoder* approach of making the prediction. We proposed a *dual encoder* approach [40] to enforce the importance of the interaction involving the peptide in the training process of the classifier. As the name suggests, we applied two encoders to process the TCR-pMHC sample. The first encoder (*ϕ*_1_) works the same way as in the single encoder approach. The second encoder (*ϕ*_1_) encodes the subgraph 𝒢 _TCR-MHC_ where all peptide nodes are removed. The encoded representations from both encoders are concatenated to make the final prediction via the MLP block. In addition to the binary cross-entropy loss ℒ _CE_, we incorporated a semantic loss ℒ _*semantic*_ (Equation (12)) to make the learned decision rule more biased toward the peptide-related interaction. Specifically, we encourage the embeddings ***g***_1_ and ***g***_2_ to have higher (lower) cosine distance for binding (non-binding) samples. Intuitively, this loss component can be thought of as a measure against over-exploiting other contexts that are not connected to peptide nodes in the residue-level graph.

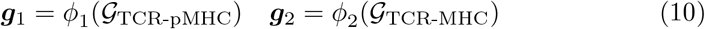

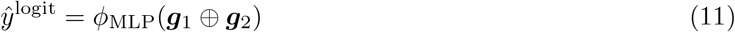

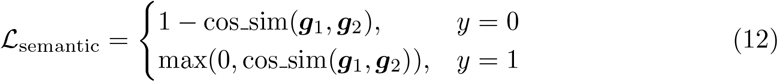

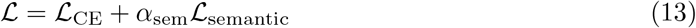

For the conditional encoding of the TCR-pMHC graph, we explored two approaches. In first approach, the TCR-MHC embedding ***g***_2_ = *ϕ*_2_(𝒢_TCR-MHC_) is used as a global feature in encoding TCR-pMHC graph according to Equation (15). The other approach utlizes ***g***_2_ (Equation (16)) to estimate the parameters of node-MLP block *ϕ*_1,node-MLP_ of *ϕ*_1_ which later encodes 𝒢_TCR-pMHC_.

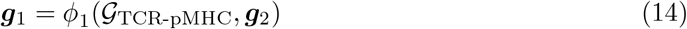

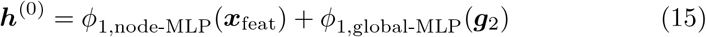

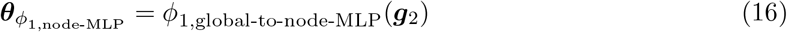

For each of the 10-fold cross-validation training (with fixed random seed for all experiments), we ran the training process with a learning rate of 0.0001 using the optimizer Adam [64] for a batch of 256 samples. We set the maximum training epochs to 350 with early stopping (patience of 25 epochs). The best model in each fold was recorded based on the ROC-AUC value on the validation split of the fold, with early stopping enabled after 50 epochs. Unless stated otherwise, the results of dual encoder approaches were from classifiers trained with *α*_sem_ = 1.

### 4.3 Training with paired preference data

For each fold’s paired preference data 𝒟^pair^ = {(𝒢_*i*_, 𝒢_*j*_)|*y*_*i*_ = 1, *y*_*j*_ = 0, Δ*E*_*ij*_ ≤ δ_th_}, we set δ_th_ = − 50 to ignore the noise in the Rosetta-estimated interaction energy. We used a preference dataset of 3000 pairs, but there are more possible pairs that can be constructed from the training split. Picking the 3000 samples based on the lowest Δ*E*_*ij*_ can lead to lowered diversity of the TCR-pMHC samples in 𝒟^pair^. Rather, we first assigned the pairs (*A*_*ij*_ = 1 instead of 0) that minimize Equation (17) (linear sum assignment problem, solved with SciPy solver [65, 66]). The remaining required samples were collected from those pairs with an unassigned TCR-pMHC sample.

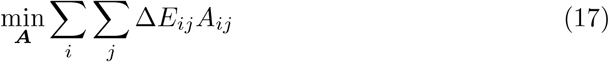

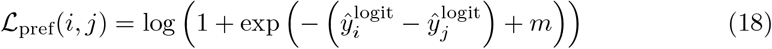

For the results shown in Section 2.6, the classifiers are periodically (at each 5th epoch starting from 51^st^ epoch) trained to minimize the ℒ_pref_ loss (Equation (18) with *m* = 3). The learning rate for this pairwise preference training was halved from the learning rate for training with binary labels. We performed early stopping after 100 epochs in this case to allow the ℒ_pref_-based training to have an impact on the selected model.

## 5 Data availability

The dataset from [28] is found at https://github.com/KavrakiLab/STAG_public. The datasets of Rosetta-relaxed and TCRmodel2 structures are shared in Zenodo (https://doi.org/10.5281/zenodo.18344673).

## 6 Code availability

We have used the codes from https://github.com/EsamTolba/TCR-CoM for computing docking and crossing angles [67] of TCR-pMHC structures. For TCRmodel2, the majority of the structures are predicted with local installation from https://github.com/piercelab/tcrmodel2 (commit 5ffb362), and the rest were done with the web-server at https://tcrmodel.ibbr.umd.edu. The codes for the experiments of this study are deposited at https://github.com/nafizabeer/TCR-pMHC-Struct-generalization.

## 7 Author contribution

**ANMNA** and **B-JY** have conceived the study. **ANMNA** designed the methodology, performed experiments, and wrote the first draft of the manuscript. **RSR, XQ, B-JY** contributed to the methodology and revised the final version of the manuscript.

## Acknowledgements

This work has been supported by the Advanced Research Projects Agency for Health (ARPA-H) Award 1AY1AX000053-01.

## Appendix A

**TCR-pMHC graph edge distribution for different holdout clusters**

**Fig. A1:**
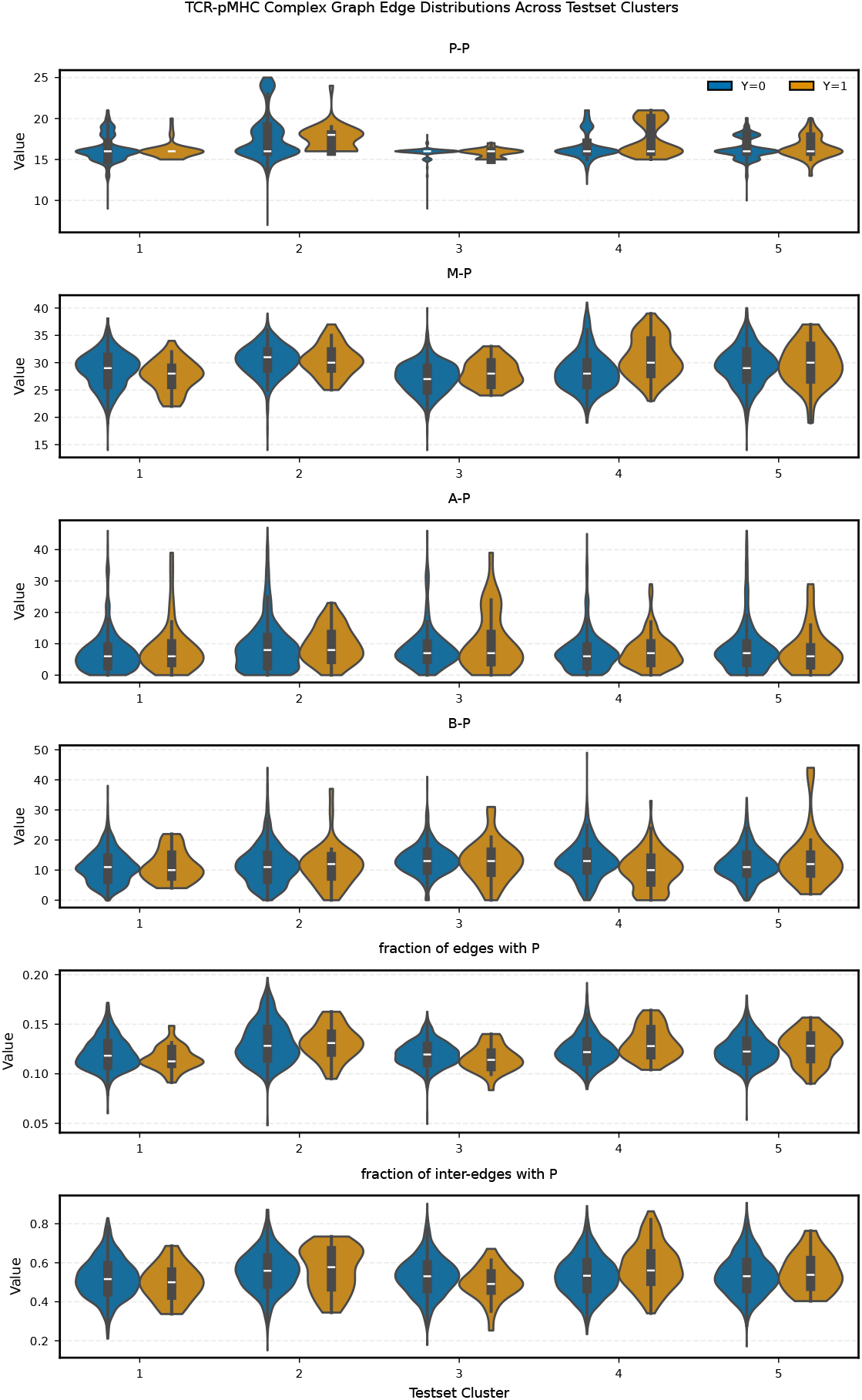
Class-specific distribution of graph edges across 5 holdout testset clusters.

Based on comparison (Figure A1) of different edge distributions across testsets, there are some unique characteristics of the testset clusters:

- The intra-chain edges for peptides show existence of multiple modes for binding samples from clusters 2, 4 and 5.
- Binding samples in clusters 4 and 5 tend to have more edges between MHC and peptide than other testset clusters.
- Cluster 5 has binding samples with more interaction between TCR-*β* and peptide.
- The TCR-pMHC interface graphs in clusters 1 and 3 has lower proportion of edges (including both inter and intra chain) connected to peptide residues compared to the other clusters. More noticeably, the cluster 4’s binding samples has larger fraction of inter-chain edges connected to peptide.

**Fig. A2:**
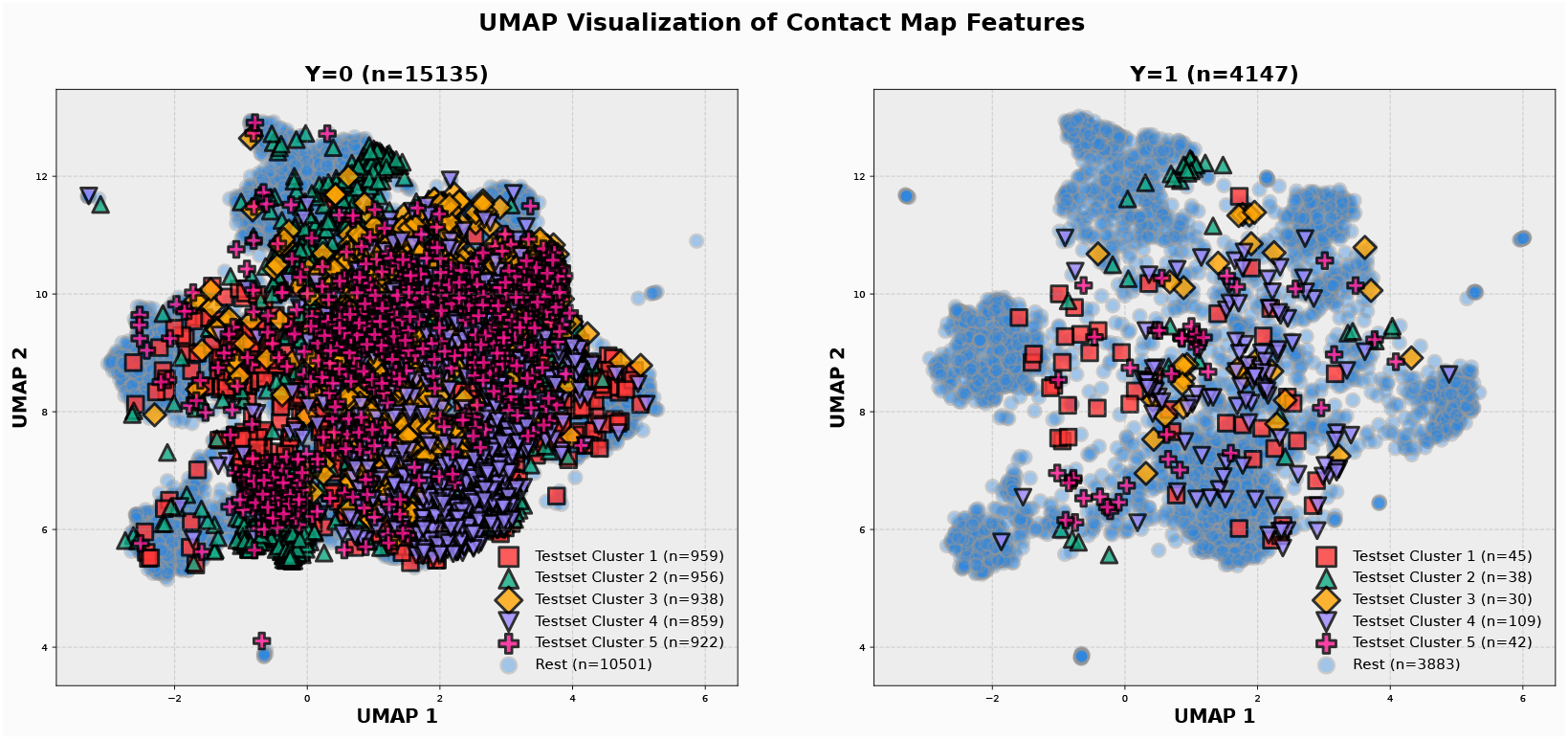
UMAP visualization of samples in *init struct* dataset. The contact map profile between group of peptide-MHC and group of *α, β* chains of TCR, i.e., 20 ×20 matrix containing normalized frequency of amino acid pairs in TCR-pMHC interface (within 8Å) is used as a feature for dimension reduction with cosine distance.

## Appendix B

**Ablation study of semantic loss weights**

In Figure B3, we show the impact of the semantic loss component on the performance of the dual encoder approach with EGNN by varying the *α*_sem_ = 0, 0.5, 1, 2. The single encoder case is also shown for reference. Without semantic loss, the dual encoder approach showed a decrease in performance, e.g., 3^rd^ quartile went down. This impact is more pronounced for clusters 4 and 5, illustrating the importance of this loss component in the robust performance of the dual encoder approach with EGNN. While increasing such loss weight led to better performance, as we see with *α*_sem_ = 2, the performance across 10 folds tends to have lower ROC-AUCs when the classifiers are over-penalized for this semantic loss.

**Fig. B3:**
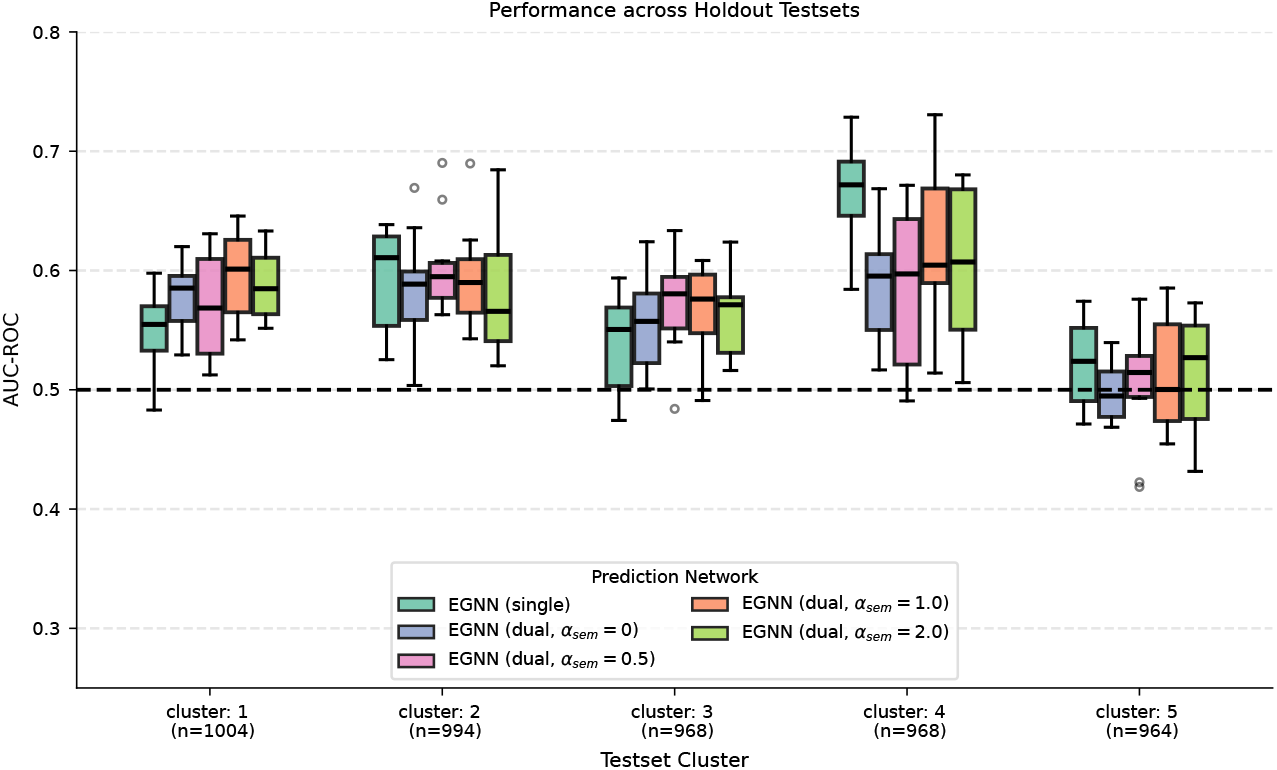
Influence of semantic loss in dual encoder approach: The single encoder and dual encoder approach with *α*_sem_ = 1.0 are same as the results reported in the main text. For the other three cases, we repeated the same 10-fold cross-validation process, followed by inference on 5 holdout testset clusters.

## Appendix C

**Aggregation of predicted probabilities: “reasonable” vs “confident” prediction**

From the 10-fold cross-validation training, we have 10 sets of learned parameters (***θ***) of the GNN-classifier. Consequently, for a test sample 𝒢_TCR-pMHC_, we have 10 predictions of binding probability, i.e., *ŷ*_*i*_ = *f* (𝒢_TCR-pMHC_; ***θ***_*i*_) with *i* ∈ {1, · · ·, 10}. To aggregate these *ŷ*_*i*_ ∈ [0, 1] into a single binding probability prediction for the test sample, we considered multiple approaches:

- ***weighted (uniform)***: equivalent to the Bayesian model averaging (BMA) technique, considering an ensemble of 10 models.

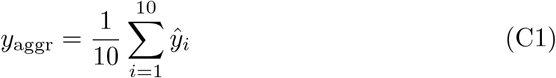
- ***best (predictive entropy)***: picking the highest confident prediction, i.e., lowest entropy 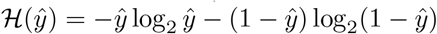

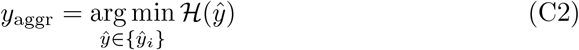
- ***weighted (predictive entropy)***: weighted average of inversely proportional to predictive entropy. *ŷ*_*i*_ with weights are inversely proportional to predictive entropy.

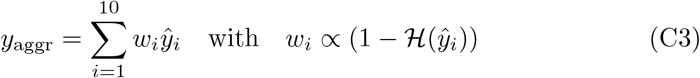

The predictive entropy only tells us about the model’s confidence in the prediction. But the model can be overconfident about the wrong prediction of binding activity. For a model trained with dataset of TCR-pMHC interface graphs, one can hypothesize that the trained model’s prediction should rely on the peptide nodes and the associated edges. Accordingly, it is intuitive to expect that a “reasonable” model will be less confident in prediction when TCR-MHC graph is presented at test time instead of the TCR-pMHC. Specifically, this prefers ℋ (*ŷ*_*i*_) to be lower than ℋ (*ŷ*_*i*,masked)_, where *ŷ*_*i*,masked_ = *f* (𝒢_TCR-MHC_; ***θ***_*i*_).

- ***best (change in predictive entropy)***:

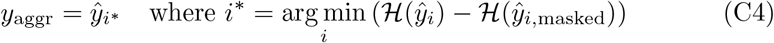
- ***weighted (change in predictive entropy)***:

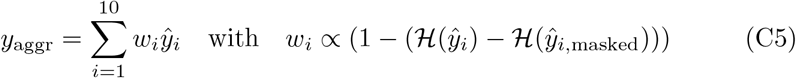

Table C1 showed the ROC-AUC values based on aggregated probabilities by applying these different aggregation approaches for the different GNN-classifiers mentioned in Section 2.1. While these aggregation approaches led to a wide range of AUC-ROC values, the use of “change in predictive entropy” seems a more intuitive choice as it offers an interpretation of “reasonable” prediction in addition to more consistent performance across different testset clusters and GNN classifiers.

**Table C1:** Performance (ROC-AUC) of different aggregation approaches. *change in predictive entropy* indicates the difference in entropy between inference with full TCR-pMHC graph vs inference with masked peptide nodes.

| Classifier | Aggregation Method | Cluster 1 | Cluster 2 | Cluster 3 | Cluster 4 | Cluster 5 |
| --- | --- | --- | --- | --- | --- | --- |
| EGNN (single) | best (change in predictive entropy) | **0.5621** | 0.6063 | **0.5819** | 0.6872 | **0.5806** |
|  | weighted (change in predictive entropy) | 0.5571 | **0.6324** | 0.5640 | 0.7105 | 0.5382 |
|  | best (predictive entropy) | 0.5372 | 0.5942 | 0.5668 | 0.6934 | 0.5604 |
|  | weighted (predictive entropy) | 0.5555 | 0.6242 | 0.5585 | 0.7022 | 0.5415 |
|  | weighted (uniform) | 0.5526 | 0.6319 | 0.5606 | **0.7109** | 0.5373 |
| EVTConv (single) | best (change in predictive entropy) | 0.6076 | **0.5764** | **0.5931** | 0.5129 | 0.4781 |
|  | weighted (change in predictive entropy) | 0.5781 | 0.5548 | 0.5728 | 0.6579 | 0.4870 |
|  | best (predictive entropy) | **0.6101** | 0.5399 | 0.5854 | 0.5356 | 0.4475 |
|  | weighted (predictive entropy) | 0.5795 | 0.5449 | 0.5615 | 0.6322 | 0.4721 |
|  | weighted (uniform) | 0.5718 | 0.5506 | 0.5766 | **0.6708** | **0.4887** |
| EGNN (dual) | best (change in predictive entropy) | 0.6231 | **0.6472** | 0.5308 | 0.6080 | 0.5372 |
|  | weighted (change in predictive entropy) | 0.6229 | 0.6217 | 0.5845 | **0.6733** | 0.5228 |
|  | best (predictive entropy) | **0.6522** | 0.6305 | **0.5990** | 0.6291 | 0.5458 |
|  | weighted (predictive entropy) | 0.6225 | 0.6274 | 0.5764 | 0.6518 | **0.5508** |
|  | weighted (uniform) | 0.6255 | 0.6205 | 0.5878 | 0.6697 | 0.5250 |
| EVTConv (dual) | best (change in predictive entropy) | **0.5910** | 0.5588 | 0.5618 | 0.5230 | 0.5263 |
|  | weighted (change in predictive entropy) | 0.5706 | 0.6001 | 0.5715 | 0.6162 | 0.5289 |
|  | best (predictive entropy) | 0.5698 | 0.6064 | **0.6155** | 0.5395 | 0.5258 |
|  | weighted (predictive entropy) | 0.5562 | **0.6168** | 0.5997 | 0.5551 | **0.5377** |
|  | weighted (uniform) | 0.5694 | 0.6011 | 0.5741 | **0.6220** | 0.5232 |

## Appendix D

**Performance with structures predicted with TCRmodel2**

**Fig. D4:**
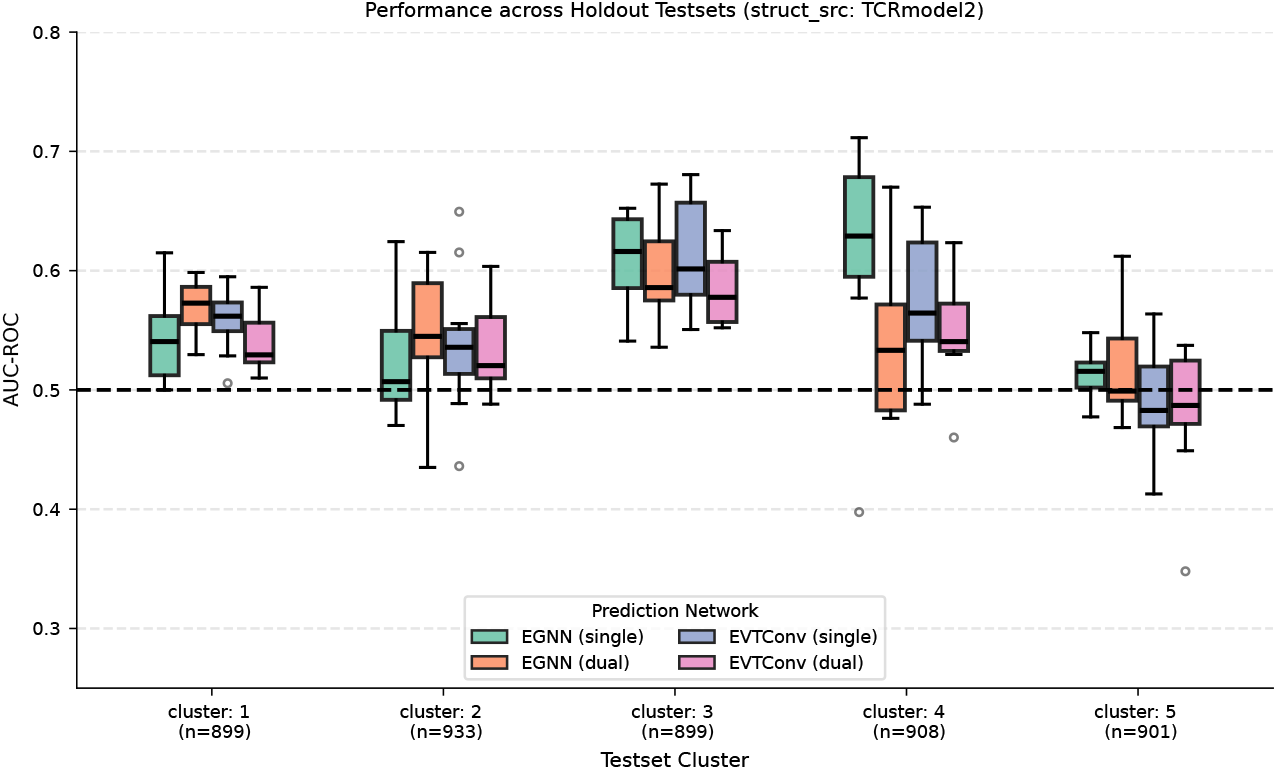
Performance of 4 different GNN-based classifiers with *tcrmodel2 struct* dataset. These four models (two types of graph encoder: EGNN and EVT-Conv, and two types of classifier architecture: single and dual encoder approach) are trained with complete TCR-pMHC graphs. The horizontal dashed line at 0.5 refers to the performance of a random classifier.

Figure D4 illustrates the generalization performance for models trained with *tcr-model2 struct* dataset. The count of testset samples is different compared to the cases of *init struct* and *relaxed struct* datasets because 987 samples are removed due to having unrealistic binding configuration, e.g., no edges between peptides and TCRs, lower model confidence. Now the trend of single vs. dual encoder (with EGNN) still exists with this dataset, similar to the case of *relaxed struct* dataset. Furthermore, cluster 5 remains a hard test cluster with this new dataset of predicted structures, making this cluster more interesting since the structural uncertainty may not be the way to tackle this challenging scenario.

## Appendix E

**Additional results on impact of HLA, data pruning and semantic loss**

**Fig. E5:**
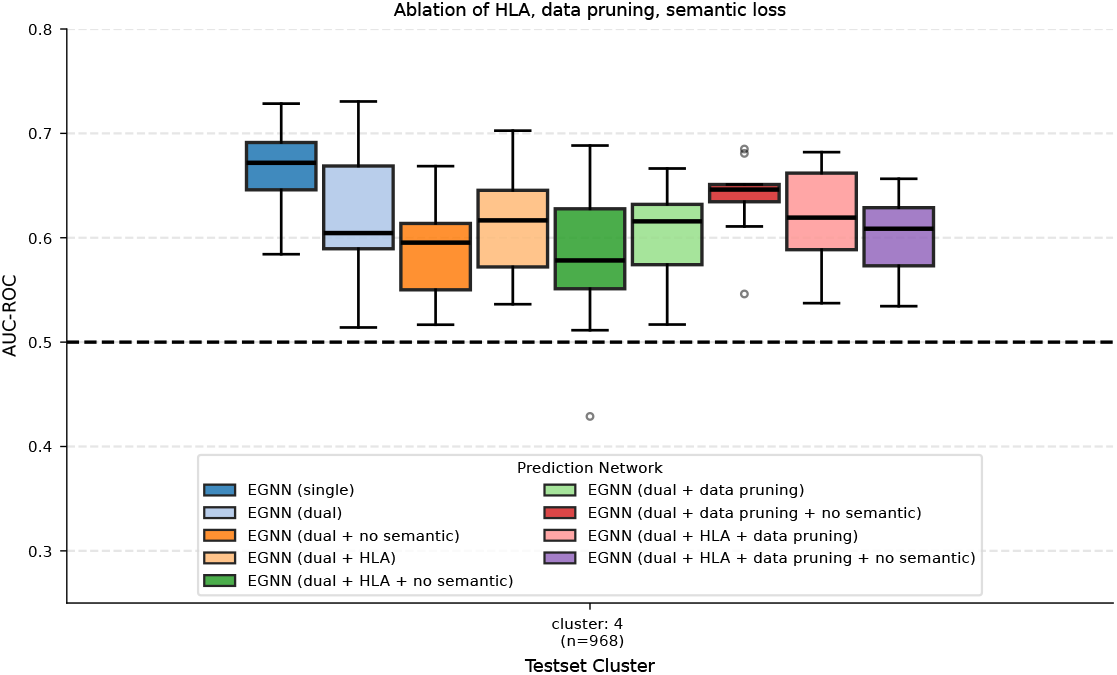
Impact of data pruning: Generalization performance in testset cluster 4 for ablation experiments of HLA, negative data pruning, and semantic loss component in dual encoder approach.

In Section 2.1, we saw that the performance dropped heavily for cluster 4 with dual approaches. We wondered whether the TCR-MHC context present in the training stage might negatively impact the performance for this cluster, since 378 out of 968 samples in this testset have MHC and TCR-*α* and *β* identical to some training set samples. We considered a negative data pruning strategy, i.e., removing the selected negative samples in the training set, which have the same (TCR, MHC) pairs as in the testset, and have positive labeled training samples with the same (TCR, MHC) pairs. Figure E5 shows the results for cluster 4 of ablation experiments of the use of HLA, data pruning, and the semantic loss component of the dual encoder approach. With the inclusion of HLA, the influence of semantic loss stays the same as in Section B, i.e., performance drop in the dual encoder with the semantic loss component. However, with only data-pruning in action, the semantic loss component became less important. Specifically, this (denoted as “*EGNN (dual + data pruning + no semantic)*” in Figure E5) moved up the median with more consistent (lower inter-quartile range) performance compared to the EGNN with the dual approach. This is more pronounced when we compare with “*EGNN (dual + no semantic)*”, highlighting the more intricate impact of the training data on the generalization performance of such structure-based classifiers.

## Appendix F

**Additional results on training with paired preference data**

**Fig. F6:**
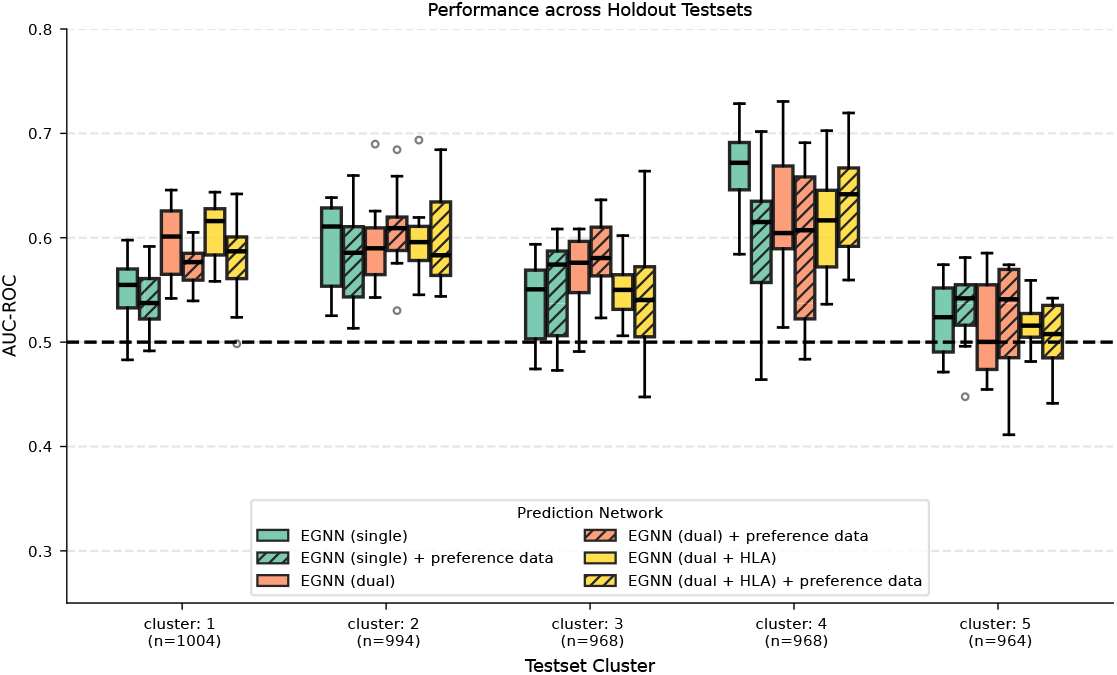
Utility of Rosetta-energy based preference dataset: Performance comparison with and without periodic training using preference dataset of 3000 pairs constructed for each fold of 10-fold cross-validation.

## References

[1] Springer, I., Tickotsky, N. & Louzoun, Y. Contribution of t cell receptor alpha and beta cdr3, mhc typing, v and j genes to peptide binding prediction. Frontiers in immunology 12, 664514 (2021).

[2] Gfeller, D. et al. Improved predictions of antigen presentation and tcr recognition with mixmhcpred2. 2 and prime2. 0 reveal potent sars-cov-2 cd8+ t-cell epitopes. Cell Systems 14, 72–83 (2023).

[3] Gao, Y. et al. Pan-peptide meta learning for t-cell receptor–antigen binding recognition. Nature Machine Intelligence 5, 236–249 (2023).

[4] Jensen, M. F. & Nielsen, M. Enhancing tcr specificity predictions by combined pan-and peptide-specific training, loss-scaling, and sequence similarity integration. elife 12, RP93934 (2024).

[5] Zhang, J., Ma, W. & Yao, H. Accurate tcr-pmhc interaction prediction using a bert-based transfer learning method. Briefings in bioinformatics 25, bbad436 (2024).

[6] Yu, C., Fang, X., Tian, S. & Liu, H. A unified cross-attention model for predicting antigen binding specificity to both hla and tcr molecules. Nature Machine Intelligence 7, 278–292 (2025).

[7] Long, X. et al. Thlanet: A deep learning framework for predicting tcr-phla binding in immunotherapy applications. PLOS Computational Biology 21, 1–20 (2025). URL 10.1371/journal.pcbi.1013050.

[8] Garboczi, D. N. et al. Structure of the complex between human t-cell receptor, viral peptide and hla-a2. Nature 384, 134–141 (1996).

[9] Gray, G. I. et al. The evolving t cell receptor recognition code: The rules are more like guidelines. Immunological Reviews 329, e13439 (2025). URL https://onlinelibrary.wiley.com/doi/abs/10.1111/imr.13439.E13439IMR-2024-116.

[10] Lin, V. et al. Tcr3d 2.0: expanding the t cell receptor structure database with new structures, tools and interactions. Nucleic Acids Research 53, D604–D608 (2024). URL 10.1093/nar/gkae840.

[11] Garcia, K. C. & Adams, E. J. How the t cell receptor sees antigen—a structural view. Cell 122, 333–336 (2005).

[12] Hummer, A. M., Schneider, C., Chinery, L. & Deane, C. M. Investigating the volume and diversity of data needed for generalizable antibody–antigen ΔΔG prediction. Nature Computational Science 5, 635–647 (2025).

[13] Fang, A. et al. Learning universal representations of intermolecular interactions with atomica. bioRxiv (2025). URL https://www.biorxiv.org/content/early/2025/07/15/2025.04.02.646906.

[14] Spoendlin, F. C. et al. Predicting the conformational flexibility of antibody and t cell receptor complementarity-determining regions. Nature Machine Intelligence 1–13 (2025).

[15] Mitra, R. et al. Geometric deep learning of protein–dna binding specificity. Nature Methods 21, 1674–1683 (2024).

[16] Motmaen, A. et al. Peptide-binding specificity prediction using fine-tuned protein structure prediction networks. Proceedings of the National Academy of Sciences 120, e2216697120 (2023). URL https://www.pnas.org/doi/abs/10.1073/pnas.2216697120.

[17] Marzella, D. F. et al. Geometric deep learning improves generalizability of mhc-bound peptide predictions. Communications Biology 7, 1661 (2024).

[18] Bradley, P. Structure-based prediction of t cell receptor: peptide-mhc interactions. Elife 12, e82813 (2023).

[19] Evans, R. et al. Protein complex prediction with alphafold-multimer. Biorxiv 2021–10 (2021).

[20] Deleuran, S. N. & Nielsen, M. Nettcr-struc, a structure driven approach for prediction of tcr-pmhc interactions. Frontiers in Immunology Volume 16 - 2025 (2025). URL https://www.frontiersin.org/journals/immunology/articles/10.3389/fimmu.2025.1616328.

[21] Basu, S. & Wallner, B. Dockq: a quality measure for protein-protein docking models. PloS one 11, e0161879 (2016).

[22] Jing, B., Eismann, S., Suriana, P., Townshend, R. J. L. & Dror, R. Learning from protein structure with geometric vector perceptrons. International Conference on Learning Representations (2021).

[23] Borrman, T., Pierce, B. G., Vreven, T., Baker, B. M. & Weng, Z. High-throughput modeling and scoring of tcr-pmhc complexes to predict cross-reactive peptides. Bioinformatics 36, 5377–5385 (2020).

[24] Culka, M. et al. Predicting specificity of tcr-pmhc interactions using machine learning and biophysical models. bioRxiv (2025). URL https://www.biorxiv.org/content/early/2025/04/07/2025.04.04.647165.

[25] Karnaukhov, V. K. et al. Structure-based prediction of t cell receptor recognition of unseen epitopes using tcren. Nature Computational Science 4, 510–521 (2024).

[26] Leaver-Fay, A. et al. in Rosetta3: An object-oriented software suite for the simulation and design of macromolecules (eds Johnson, M.L. & Brand, L.) Computer Methods, Part C, Vol. 487 of Methods in Enzymology 545–574 (Academic Press, 2011).

[27] Huang, J. & MacKerell Jr, A. D. Charmm36 all-atom additive protein force field: Validation based on comparison to nmr data. Journal of computational chemistry 34, 2135–2145 (2013).

[28] Slone, J. K., Conev, A., Rigo, M. M., Reuben, A. & Kavraki, L. E. Tcr-pmhc binding specificity prediction from structure using graph neural networks. IEEE Transactions on Computational Biology and Bioinformatics (2025).

[29] Slone, J. K. et al. Stag-llm: Predicting tcr-phla binding with protein language models and computationally generated 3d structures. Computational and Structural Biotechnology Journal 27, 3885–3896 (2025). URL https://www.sciencedirect.com/science/article/pii/S2001037025003642.

[30] Lin, Z. et al. Language models of protein sequences at the scale of evolution enable accurate structure prediction. bioRxiv (2022).

[31] Lin, Z. et al. Evolutionary-scale prediction of atomic-level protein structure with a language model. Science 379, 1123–1130 (2023).

[32] Wang, Z. et al. Lm-gvp: an extensible sequence and structure informed deep learning framework for protein property prediction. Scientific reports 12, 6832 (2022).

[33] Li, X., Sun, C., Huang, W., Wang, Y. & Ma, B. Sagetcr: a structure-based model integrating residue- and atom-level representations for enhanced tcr-pmhc binding prediction. Briefings in Bioinformatics 26, bbaf496 (2025).

[34] Abramson, J. et al. Accurate structure prediction of biomolecular interactions with alphafold 3. Nature 630, 493–500 (2024).

[35] Yin, R. et al. Tcrmodel2: high-resolution modeling of t cell receptor recognition using deep learning. Nucleic Acids Research 51, W569–W576 (2023). URL 10.1093/nar/gkad356.

[36] Passaro, S. et al. Boltz-2: Towards accurate and efficient binding affinity prediction. BioRxiv 2025–06 (2025).

[37] McMaster, B., Thorpe, C., Ogg, G., Deane, C. M. & Koohy, H. Can alphafold’s breakthrough in protein structure help decode the fundamental principles of adaptive cellular immunity? Nature Methods 21, 766–776 (2024).

[38] Grazioli, F. et al. On tcr binding predictors failing to generalize to unseen peptides. Frontiers in immunology 13, 1014256 (2022).

[39] Castorina, L. V., Grazioli, F., Machart, P., Mösch, A. & Errica, F. Assessing the generalization capabilities of tcr binding predictors via peptide distance analysis. PLOS ONE 20, 1–15 (2025). URL 10.1371/journal.pone.0324011.

[40] Abeer, A. N. M. N., Qian, X. & Yoon, B.-J. A Dual Graph Encoder Approach to Structure-Based TCR-pMHC Binding Prediction (Association for Computing Machinery, New York, NY, USA, 2025). URL 10.1145/3765612.3767776.

[41] Satorras, V. G., Hoogeboom, E. & Welling, M. Meila, M. & Zhang, T. (eds) E(n) equivariant graph neural networks. (eds Meila, M. & Zhang, T.) Proceedings of the 38th International Conference on Machine Learning, Vol. 139 of Proceedings of Machine Learning Research, 9323–9332 (PMLR, 2021). URL https://proceedings.mlr.press/v139/satorras21a.html.

[42] Delgado, J. et al. Foldx force field revisited, an improved version. Bioinformatics 41, btaf064 (2025). URL 10.1093/bioinformatics/btaf064.

[43] Leman, J. K. et al. Macromolecular modeling and design in rosetta: recent methods and frameworks. Nat. Methods 17, 665–680 (2020).

[44] Zareie, P. et al. Canonical t cell receptor docking on peptide–mhc is essential for t cell signaling. Science 372, eabe9124 (2021).

[45] Qin, R. et al. Tcr catch bonds nonlinearly control cd8 cooperation to shape t cell specificity. Cell Research 1–19 (2025).

[46] Resink, T. et al. From structure to immunogenicity: Decoding correlated dynamics at the peptide mhc interface to understand tcr recognition. bioRxiv (2025). URL https://www.biorxiv.org/content/early/2025/05/23/2025.05.18.654692.

[47] Quast, N. P. et al. T-cell receptor structures and predictive models reveal comparable alpha and beta chain structural diversity despite differing genetic complexity. Communications Biology 8, 362 (2025).

[48] Cagiada, M., Spoendlin, F. C., Ifashe, K. & Deane, C. M. Uncovering the flexibility of cdr loops in antibodies and tcrs through large-scale molecular dynamics. bioRxiv 2025–11 (2025).

[49] Jing, B., Berger, B. & Jaakkola, T. Salakhutdinov, R. et al. (eds) AlphaFold meets flow matching for generating protein ensembles. (eds Salakhutdinov, R. et al.) Proceedings of the 41st International Conference on Machine Learning, Vol. 235 of Proceedings of Machine Learning Research, 22277–22303 (PMLR, 2024).

[50] Stratiichuk, R. et al. Sampling and ranking of protein conformations using machine learning techniques do not improve the quality of rigid protein–protein docking. Journal of Chemical Information and Modeling 65, 10167–10179 (2025).

[51] Riley, T. P. et al. T cell receptor cross-reactivity expanded by dramatic peptide– mhc adaptability. Nature chemical biology 14, 934–942 (2018).

[52] Visani, G. M. et al. T-cell receptor specificity landscape revealed through de novo peptide design. bioRxiv (2025). URL https://www.biorxiv.org/content/early/2025/09/04/2025.02.28.640903.

[53] Liu, B. et al. Design of high-specificity binders for peptide–mhc-i complexes. Science 389, 386–391 (2025). URL https://www.science.org/doi/abs/10.1126/science.adv0185.

[54] Householder, K. D. et al. De novo design and structure of a peptide–centric tcr mimic binding module. Science 389, 375–379 (2025). URL https://www.science.org/doi/abs/10.1126/science.adv3813.

[55] Tickotsky, N., Sagiv, T., Prilusky, J., Shifrut, E. & Friedman, N. Mcpas-tcr: a manually curated catalogue of pathology-associated t cell receptor sequences. Bioinformatics 33, 2924–2929 (2017).

[56] Shugay, M. et al. Vdjdb: a curated database of t-cell receptor sequences with known antigen specificity. Nucleic acids research 46, D419–D427 (2018).

[57] Vita, R. et al. The immune epitope database (iedb) 3.0. Nucleic acids research 43, D405–D412 (2015).

[58] 10x Genomics. A new way of exploring immunity–linking highly multiplexed antigen recognition to immune repertoire and phenotype. Tech. rep (2019).

[59] Jensen, K. K. et al. Tcrpmhcmodels: Structural modelling of tcr-pmhc class i complexes. Scientific reports 9, 14530 (2019).

[60] Lusiany, T. et al. Structural Modeling of Adaptive Immune Responses to Infection, 283–294 (Springer US, New York, NY, 2023). URL 10.1007/978-1-0716-2609-215.

[61] Heather, J. M. et al. Stitchr: stitching coding tcr nucleotide sequences from v/j/cdr3 information. Nucleic acids research 50, e68–e68 (2022).

[62] Atchley, W. R., Zhao, J., Fernandes, A. D. & Drüke, T. Solving the protein sequence metric problem. Proceedings of the National Academy of Sciences 102, 6395–6400 (2005).

[63] Kidera, A., Konishi, Y., Oka, M., Ooi, T. & Scheraga, H. A. Statistical analysis of the physical properties of the 20 naturally occurring amino acids. Journal of Protein Chemistry 4, 23–55 (1985).

[64] Kingma, D. P. & Ba, J. Adam: A method for stochastic optimization. ICLR (2015).

[65] Virtanen, P. et al. SciPy 1.0: Fundamental Algorithms for Scientific Computing in Python. Nature Methods 17, 261–272 (2020).

[66] Crouse, D. F. On implementing 2d rectangular assignment algorithms. IEEE Transactions on Aerospace and Electronic Systems 52, 1679–1696 (2016).

[67] Singh, N. K. et al. Geometrical characterization of t cell receptor binding modes reveals class-specific binding to maximize access to antigen. Proteins: Structure, Function, and Bioinformatics 88, 503–513 (2020). URL https://onlinelibrary.wiley.com/doi/abs/10.1002/prot.25829.

